# TMS SMART – Scalp Mapping of Annoyance Ratings and Twitches caused by Transcranial Magnetic Stimulation

**DOI:** 10.1101/191320

**Authors:** Lotte Meteyard, Nicholas Holmes

## Abstract

The magnetic pulse generated during Transcranial magnetic stimulation [TMS] also stimulates cutaneous nerves and muscle fibres, with the most commonly reported side effect being muscle twitches and sometimes painful sensations. These sensations affect behaviour during experimental tasks, presenting a potential confound for ‘online’ single-pulse TMS studies. Our objective was to systematically map the degree of disturbance (ratings of annoyance, pain, and muscle twitches) caused by TMS at 43 locations across the scalp. Ten participants provided ratings whilst completing a choice reaction time task, and ten participants provided ratings whilst completing a ‘flanker’ reaction time task. TMS over frontal and inferior regions resulted in the highest ratings of annoyance, pain, and muscle twitches caused by TMS. In separate analyses we predicted the difference in reaction times (RT) under TMS by scalp location and subjective ratings. Frontal and inferior scalp locations showed the greatest cost to RTs under TMS (i.e., slowing), with midline sites showing no or minimal slowing. Increases in subjective ratings of disturbance predicted longer RTs under TMS. Critically, ratings were a better predictor of the cost of TMS than scalp location or scalp-to-cortex distance, and the more difficult ‘flanker’ task showed a greater effect of subjective disturbance. The peripheral sensations and discomfort caused by TMS pulses significantly and systematically influence RTs during single-pulse, online TMS experiments. We provide the data as an online resource (http://www.tms-smart.info) so that researchers can select control sites that account for the level of general interference in task performance caused by online single-pulse TMS.

## 1.0 Introduction

A transcranial magnetic stimulation (TMS) machine produces a rapidly-changing magnetic field which, when positioned on the scalp, can be used to disrupt function temporarily in a specific brain area (Sparing & Mottaghy, 2008; Wagner, Valero-Cabre, & Pascual-Leone, 2007; Walsh & Cowey, 2000). This provides a non-invasive means to draw inferences about the contribution of specific brain regions to a given behavioural task. (Bolognini & Ro, 2010; Ruff et al., 2009).

TMS causes peripheral auditory and cutaneous sensations (Bestmann et al., 2005; Borckardt et al., 2006; Nikouline, Ruohonen, & Ilmoniemi, 1999; Starck et al., 1996; Wasserman, 1998). Tension headaches and scalp discomfort are frequently reported side-effects of undergoing TMS (Anderson et al., 2006; Benninger et al., 2011; Maizey et al., 2013; O’Reardon, and et al., 2007; VonLoh et al., 2013).

These peripheral sensations differ by scalp location. TMS is more disturbing over prefrontal cortex than over parietal cortex (Abler et al., 2005). Crying has been reported as an adverse effect for participants in speech studies that stimulated left prefrontal areas (Wasserman, 1998). Greater discomfort over frontal regions of the scalp is likely due to a greater density of muscles and nerves (Maizey et al., 2013; Wassermann, 1998; Machii et al., 2006; Loo et al., 2008). Discomfort is related to the intensity and frequency of stimulation but even single-pulse TMS can cause notable discomfort, especially with participants naïve to TMS (Maizey et al., 2013). Notably, ratings of scalp discomfort caused by TMS positively correlate with the number of errors made on a delayed match to sample task (Abler et al., 2005).

Appropriate control conditions are needed to ensure that results are due to the changes in brain activity caused by TMS, not as a consequence of peripheral effects (Arana et al., 2008; Duecker & Sack, 2013 & 2015; Sandrini et al., 2011). There are several options for control conditions in TMS: the use of additional behavioural tasks or conditions, a no TMS condition (e.g., Tamè & Holmes, 2016), the use of sham TMS and application of TMS to a control location on the scalp.

Duecker & Sack (2015) provided a review of sham TMS, for which a number of conditions and devices have been evaluated and developed (e.g., Arana et al., 2008; Rossi et al., 2007; Sommer et al., 2006). Sham TMS is designed to mimic the sound and sometimes the peripheral sensations of TMS, without changing cortical activity (Borckardt et al., 2008; Duecker & Sack, 2015), and it is possible to create a sham condition under which participants cannot reliably judge whether they received sham or real rTMS (Duecker & Sack, 2015; Herwig et al., 2010). However, sham TMS has to deliver high intensity cutaneous sensations (70% of participant’s maximum pain tolerance) to be matched to real TMS (Arana et al, 2008).

Control locations for TMS should attempt to control for peripheral sensations, yet to our knowledge, this control is not often implemented. Ideally, a selected control site should give the same level of comfort or discomfort as the experimental site. The vertex (Cz, at the top of the head) is often used, under the assumption that the brain tissue is deeper under the scalp, attenuating cortical stimulation (e.g., Kalbe et al., 2010; Sandrini et al., 2011; Silvanto et al., 2008; Whitney et al., 2010; Vetter et al., 2015; but see Fox et al., 1997; Jung et al., 2016; Loo et al., 2000). It is not clear that this location controls well for cutaneous sensations and discomfort, particularly when the experimental site is frontal and known to be more painful (Wasserman, 1998).

The best-practice in TMS studies may be to have more than one control condition, with multiple stimulation sites or stimulation time points (Duecker & Sack, 2015; Miall et al,. 2008; Tamè & Holmes, 2016) and a control task (Sandrini et al., 2011). However, the selection of control conditions remains sub-optimal.

The aim of the study reported here is to map the subjective discomfort of TMS across the scalp. Rather than comparing a small number of sites, we aimed for coverage of the entire scalp, in order to create a detailed map of the frequency of muscle twitches, and participant ratings of annoyance and pain (extending Abler et al, 2005; Arana et al., 2008; Loo et al., 2000). We used two different tasks to test whether task difficulty would interact with the application of. That is, does greater discomfort under TMS affect a difficult task more than an easier task? Together with an accompanying website (http://www.tms-smart.info), this data provides a resource that researchers can use to control for peripheral sensations caused by TMS.

## 2.0 Method

### 2.1 Participants

Twenty individuals took part. Ten completed a choice-RT (CRT) ask and ten completed a ‘flanker’ task. All participants provided written informed consent and underwent screening in line with the TMS Rules of Operation at the Centre for Integrative Neuroscience and Nuerodynamics, University of Reading. The study received a favourable opinion for conduct from the University of Reading Research Ethics Committee (UREC Project 13/03).

For the choice RT task, mean age was 31.6 years (SD=5.5), 6 were female and all were right handed by self-report. Mean resting motor threshold, examined by visual inspection of muscle twitches in the hand, was 51.9% (SD=7.7%) of maximum stimulator output. For the flanker task, mean age was 25.2 years (SD=3.5 years), 6 were female and 8 were right handed by self-report. Their mean resting motor threshold was 48.5% (SD=6.4%) of maximum stimulator output (Rossini et al., 1994). All participants completed the ratings task. No participants or other data were excluded from the final dataset prior to analysis.

### 2.2 Scalp locations

We aimed for coverage of the entire scalp. To begin, we transferred locations from a 32 channel EEG cap onto a Lycra skull cap (see Herwig et al., 2003 and Homan et al., 1987 for the 10:20 system being used position the TMS coil). The 32 locations were marked using BrainSight Frameless Stereotaxy (Rogue Resolutions, Cardiff, Wales), and registered to a T1 structural scan (author NPH). This registration made clear that the occipital, temporal, and cerebellar areas were not fully covered, so we added 5 sites per hemisphere and one midline site. This gave a total of 24 scalp locations in each hemisphere (5 in the midline, and 19 covering each hemisphere; Figure 1). For occipital sites, we used 6 cm distance measures as a rough guide from point to point marked on the cap using the 10:20 system. For the midline we took a line from Cz to Oz (our location: Inion). We also took lines from Pz to O2/O3 (our location: Lateral Inion 1 & 2), P3 to P7/P8 (our location: Lateral Occipital 1 & 2). For other sites, we used the registration data available in BrainSight to add locations approximately corresponding to the following: Lateral-Occipital (3 & 4), Temporal-Parietal Junction (TPJ 1 & 2) and the Anterior Temporal Lobes (ATL 1 & 2; see Figure 1).

**Figure 1:**
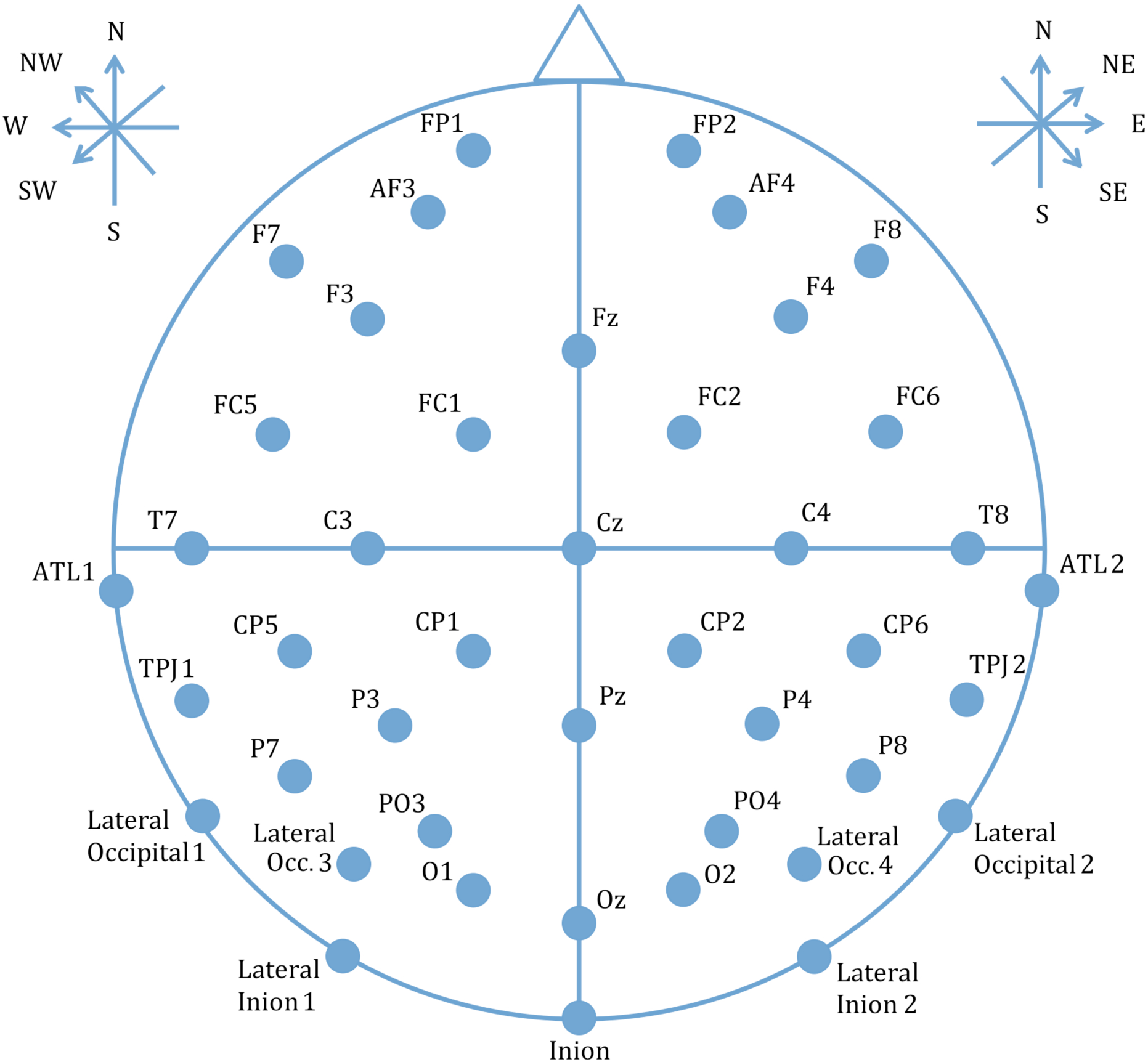
Locations of cortical sites stimulated with TMS and axes of stimulation. Legend/Key: Scalp locations on the Lycra cap were assigned numbers to support randomisation and blinding during the experiment. For each participant Cz was aligned to their vertex and this alignment was checked periodically during testing.

### 2.3 Coil orientations

To measure the influence of coil orientation we used a cardinal reference system with North taken as the direction faced by the participant. This was preferable to any allocentric (e.g., superior, inferior, anterior, posterior) or neuraxis-centred (e.g., ventral, dorsal, rostral, caudal) reference system, since coil orientation did not change as the coil moved along the scalp. ‘South’ was always with the handle pointing along the surface of the scalp following ‘lines of longitude’ towards the inion; ‘East’ was always with the handle pointing along the surface of the scalp following similar lines of longitude (not latitude) towards the right ear. Each location was stimulated biphasically using four coil orientations that mapped 45 degree rotations around a circle. Orientations were North-South (along the midline), North East-South West (45° going clockwise), East-West (along the ear-to-ear axis, 90°), and North West-South East (135°, see Figure 1). The TMS coil handle was aligned to the above axes, usually oriented towards the back of the participant’s head, and away from the face. It was rotated to lie in the opposite direction along the axis if this position was not possible (e.g., if the shoulder was in the way for inferior and lateral sites).

### 2.4 TMS intensity

We kept the intensity constant at 50% of the maximum stimulator output. This is the approximate mean intensity for eliciting MEPs and visible muscle contractions with this equipment in our laboratory. Researchers using TMS often change the intensity of stimulation on a participant-by-participant basis, for example, by setting the intensity to a fixed proportion of the resting (RMT) or active motor threshold (AMT). This approach is appropriate when stimulating over motor cortex in order to affect or elicit motor-evoked potentials in a contralateral muscle. A similar approach may be useful for eliciting phosphenes by stimulating over occipital cortex (Siniatchkin et al., 2011). However, much less is known about the relationship between TMS intensity and assumed cortical activation for all other brain areas (e.g. Peurala et al., 2008). We chose 50% intensity to provide a realistic level of stimulation (at approximate RMT) within the typical range for a TMS study, and therefore reflecting a typical level of discomfort. This level of stimulation is likely to have stimulated the cortex of many, if not all of the sites used in our experiment. However, because of differences in scalp to brain distance between participants and sites (Table 1), the amount of cortical stimulation likely differed across sites. We had two options for setting TMS intensity: 1) adjusted for each participant and site, or 2) kept constant across participants and sites.

**Table 1:** Least squares estimated means^1^ for the effect of TMS on reaction times (ms) by scalp location with approximate MNI co-ordinates (mean of left and right hemispheres, with absolute X coordinates) and scalp-cortex distance (mm)^2^

| **Location** | Estimated mean (ms) | t-value<sup>1</sup> | Approximate MNI | Mean+/-SD scalp-cortex distance (mm) |
| --- | --- | --- | --- | --- |
| CZ | -1.96 | -0.14 | (6,-13,99) | 17.7 (16.7,18.6) |
| FZ | 13.96 | 1.03 | (9,44,73) | 15.0 (14.0,16.0) |
| PZ | -16.83 | -1.24 | (4,-68,95) | 21.2 (19.7,22.6) |
| OZ | 11.08 | -0.82 | (17,-102,42) | 13.6 (12.5,14.6) |
| FP | 45.83 | 3.37* | (30,77,25) | 13.1 (12.1,14.2) |
| AF | 27.51 | 2.03 | (37,64,41) | 14.4 (13.3,15.4) |
| F3&4 | 4.60 | 0.34 | (41,32,66) | 14.2 (13.3,14.9) |
| F7&8 | 71.27 | 5.26* | (71,27,21) | 13.4 (12.3,14.4) |
| FC1&2 | 6.28 | 0.46 | (28,16,86) | 15.1 (14.1,16.0) |
| FC5&6 | 48.48 | 3.58* | (67,5,54) | 13.8 (12.9,14.7) |
| C3&4 | 2.92 | 0.21 | (47,-18,86) | 14.4 (13.3,15.4) |
| T7&8 | 56.49 | 4.15* | (78,-26,39) | 11.6 (10.6,12.5) |
| CP1&2 | -10.46 | -0.77 | (28,-48,96) | 17.1 (16.0,18.1) |
| CP5&6 | 5.44 | 0.40 | (65,-47,68) | 16.3 (15.2,17.5) |
| P3&4 | -6.25 | -0.46 | (39,-70,83) | 18.9 (17.7,20.0) |
| P7&8 | 16.89 | 1.25 | (60,-77,45) | 13.6 (12.8,14.3) |
| PO3&4 | -22.88 | -1.69* | (37,-87,66) | 16.0 (14.9,17.1) |
| O1&2 | -9.57 | -0.71 | (33,-98,43) | 13.4 (12.4,14.4) |
| Inion | 35.38 | 2.61 | (20,-115,-41) | 21.4 (19.3,23.8) |
| Lateral-inion | 47.74 | 3.52* | (51,-98,-46) | 17.4 (15.9,19.1) |
| Lateral-Occipital 1&2 | 65.33 | 4.80* | (58,-66,-42) | 15.3 (14.1,16.7) |
| Lateral Occipital 3&4 | 9.93 | 0.73 | (59,-101,-11) | 11.8 (10.9,12.6) |
| ~TPJ | 31.99 | 2.35 | (47,-55,17) | 11.7 (11.1,12.4) |
| ~ATL | 74.56 | 5.50* | (63,-14,-2) | 11.4 (10.7,12.1) |

<sup>1</sup> Least squares means testing whether RT difference (TMS-No TMS) for each scalp location is significantly different to zero. Positive numbers indicate slower reaction times under TMS, negative numbers indicate faster reaction times under TMS. Least squares means have been calculated for a model that includes Scalp Location X Hemisphere, Task and Trial as predictors.
<sup>2</sup> Measured in a separate group of 20 participants for whom we had an MRI scan available. The brain and TMS locations used in the main experiment was registered to each of 19 other brain scans, and mean scalp-cortex distance was estimated using FLSView.

We could have attempted to set the TMS intensity to obtain approximately the same level of cortical stimulation across all participants and sites. Using MEPs as an output variable, systematically mapping over the motor cortex to find the site and coil orientation that induces MEPs at the lowest intensity, then found the RMT and AMT for each participant, then decreased the intensity so that no effect of TMS was visible on the electromyograph (EMG). Using this ineffectual TMS intensity, we could then have applied a scalp to brain distance adjustment (Stokes et al., 2007) to adjust the TMS intensity for all remaining TMS sites. This method would have the advantage of being less likely to have stimulated the cortex, but it would have likely resulted in TMS intensities of 20-30% of the maximum stimulator output, much lower than typically used in experiments. Most importantly, this method of TMS intensity adjustment per participant and site would have confounded the purpose of the experiment: to measure the relative annoyance of TMS between scalp sites at levels of stimulation typically used in experiments. In addition, different TMS intensities at different scalp locations would render the ratings of annoyance, pain and twitches meaningless for further analysis. In order to systematically map the difference in subjective discomfort across scalp sites, TMS intensity needed to be constant whilst we differed scalp location.

Thus, we maintained TMS intensity constant, at 50% of maximum stimulator output. Ours is a study of the scalp, not of the brain. We accept, of course, that stimulation of the underlying cortex was very likely for many, if not all scalp locations and participants. Nevertheless, it is parsimonious to assume that our results relate best to regions of the scalp, targeted with constant-intensity TMS, rather than to regions of the brain, not targeted and with variable-intensity TMS. We also completed post-hoc analyses to control for scalp-to-brain distance (see below).

### 2.5 Tasks and stimuli

All stimuli were presented using Matlab and Psychtoolbox 3 (Brainard, 1997) in black text on a white background, font size 20, screen resolution 1920 × 1080 pixels. Stimuli were presented using a CRT monitor with 75 Hz refresh rate.

#### 2.5.1 Choice reaction time task

This task was a basic spatial-compatibility, choice RT task designed to be simpler than the flanker task (below). Stimuli were the left and right chevrons (< and >). A fixation cross was presented centrally. Three chevrons appeared on the left or right of the fixation cross, displaced 100 pixels horizontally from the central fixation cross. These could be left or right facing, as above. Therefore, there were two congruent conditions (<<< +, + >>>) and two incongruent conditions (>>> +, + <<<). Participants were instructed to press C with one finger if the chevron appeared on the left, and N with another finger if it appeared on the right. These are spatially-compatible responses as the C key is on the left and the N key on the right of the keyboard. Speed and accuracy were both encouraged. When the data were inspected, three participants had responded to arrow direction rather than the side of the screen on which the arrow appeared. As this still constituted a choice RT task with spatial compatibility (the congruence conditions were still valid), as the behavioural task was not the primary purpose of the present report, and as their data were comparable to other participants’ (mean RT, proportion correct, and size of congruency effect, see Appendix A), these three participants’ data were included in the final analysis. Removing these data points and re-analysing did not change the conclusions drawn.

#### 2.5.2 Flanker task (Eriksen & Eriksen, 1974)

Stimuli were the left and right chevrons (< and >). Three chevrons were presented centrally, and faced left (<<<) or right (>>>). Flanker chevrons facing left or right were presented either side, displaced 100 pixels horizontally from the centre of the screen. Therefore, there were two congruent conditions (<<< <<< <<<, >>> >>> >>>) and two incongruent conditions (<<< >>> <<<, >>> <<< >>>).Participants were instructed to press the C key with one finger if the central chevrons pointed left, and N with another finger if they pointed right. Speed and accuracy were both encouraged.

#### 2.5.3 Ratings Task

Participants were asked to rate the annoyance, pain, and peripheral muscle twitches in the head, face and neck caused by TMS. Each was rated on a scale of 0 to 10, with 0 being the minimum and 10 the maximum. For annoyance (“How annoying were the TMS pulses?”), anchor points (printed on a sheet of paper for reference) were “not at all annoying” (0) to “highly distracted and unable to complete the task” (10). For pain (“If painful, how intense was the pain from the TMS pulses?”), anchors were “no pain at all” (0) to “the worst pain they could tolerate during the experiment” (10). We specified pain intensity, so that participants would focus on the sensory aspects of the pain (intensity, quality) rather than their emotional response (i.e., pain unpleasantness) that would already be partly captured by the rating of annoyance (Melzack & Casey, 1968). For twitches (“If there were any twitches, how strong were the muscle twitches from the TMS pulse?”), anchors were “no twitches” (0) to “a very strong cramp” (10). In addition to the subjective ratings provided by the participants, a second experimenter recorded whether any visible twitches were observed in the face, head or neck that were coincident with the TMS pulse (0 to 5 twitches observed in each block). To facilitate this, a mirror was angled in front of the participant so the researcher could sit or stand behind them and observe. The participant was not able to observe themself in the mirror.

### 2.6 Design and procedure

The Lycra cap with scalp locations was fitted to the participant with Cz aligned to their vertex, and periodically checked during the experiment. Resting motor threshold was first established for each participant (Rossini et al., 1994). We identified the approximate location of M1 by measuring 5cm lateral and 1cm anterior from vertex for the hemisphere to be tested (right or left). Participants rested their ipsilateral hand on the desk, clearly visible to the experimenters. Test TMS pulses were delivered around this location until a visible twitch was seen in the thumb, index, or middle fingers (typically originating from the 1^st^ or 2^nd^ dorsal interosseous, adductor pollicis, or opponens pollicis muscles). Stimulator intensity was varied until a visible twitch was seen on 5 out of 10 consecutive pulses delivered at approximately 0.2Hz. This stimulator intensity was recorded as the resting motor threshold (RMT).

For the behavioural experiment, stimulation strength was set at 50% of the maximum stimulator output (biphasic). As confirmed by our data, this level was selected to be approximately at RMT. Single-pulse TMS studies typically stimulate at 110-140% RMT. However, since RMT assessed visually (as we did) is likely to be 11% higher than RMT assessed with EMG (Westin et al., 2014), this intensity of stimulation was likely above threshold for stimulating cortex. One hemisphere per participant was stimulated at all scalp locations during the experiment. The order of testing Scalp Location was fully randomised for each participant. Coil orientation was nested within location and then randomised (i.e., a randomly selected permutation of all possible orders of the four coil orientations was selected once location had been selected). Half of the participants had their left, and half their right hemisphere stimulated. This between-subjects manipulation of hemisphere was a pragmatic decision that allowed data collection to be completed in one two-hour session (partly to meet the requirements of our local ethics committee). To account for any potential confound from this manipulation, data analyses included Hemisphere and the interaction between Hemisphere and Scalp Location as predictors of interest.

Participants provided button press responses using the hand ipsilateral to the TMS, to minimise any chance of cortical stimulation from the TMS influencing hand motor control and response times. Single pulse TMS was delivered on each trial using a figure-of-eight coil.

Each task had identical timings and trial and block structure. Participants completed 60 practice trials with no TMS (coil disabled, away from head; 3 blocks of 20 trials, with each block split into 4 mini-blocks of 5 trials to reflect the actual experiment). They were then introduced to the rating task and response scales. To explain the rating procedure, participants were asked to remember the TMS pulses delivered during the RMT procedure and consider the sensations (annoyance, pain, and twitches) that they may have experienced. Thus, these pulses acted as reference pulses for participants’ ratings of annoyance, pain, and twitches. During the experiment, participants completed 5 experimental trials for each combination of coil orientation and scalp location. The experiment was structured so that a location was randomly selected, and then each of 4 randomly-ordered coil orientations was completed for that location (4 blocks of 5 trials for each location; 20 trials per location). In each block of 5 trials, a fixation cross appeared for a randomly selected duration between 500 and 1000ms, followed by the imperative stimulus. A single TMS pulse was delivered at the same time as the imperative stimulus appeared on screen. Stimuli appeared on screen until the participant responded or until 2 seconds, whichever was sooner. There was an inter-trial-interval of 2 seconds. In each block of 5 trials, participants saw one of each trial type (congruent left, incongruent left, congruent right, incongruent right) and a randomly selected 5^th^ trial. After completing the 5 trials of the task, they were asked to give ratings for the annoyance, pain, and twitches caused by the preceding 5 TMS pulses. Participants were asked to provide ratings after 5 trials to allow them to complete the behavioural task whilst experiencing more than one pulse at that location and with that coil orientation. The total number of trials per participant was 480 (5 trials x 4 orientations x 24 locations). In between each block of 5 trials, participants were able to take a break for as long as they wished.

### 2.7 Analysis

Ratings data were taken from all 20 participants and averaged across participants to give values for each coil orientation at each scalp location. Mean ratings (annoyance, pain, twitches) and measurement of observed visible twitches were strongly positively correlated with each other (all correlation coefficients>=0.85). This precludes them being used as separate fixed effect predictors due to collinearity. The self-reported intensity of twitches correlated the most strongly with all other variables (all correlation coefficients>=0.9), and was therefore used in further analyses as a fixed effect predictor.

RTs were taken from correct trials only. We used RTs from correct practice trials for baseline RT (the first 20 trials were discarded as true practice trials - we assumed that RT would be stable after these initial trials). The mean practice RT was taken for each trial type (congruent left, incongruent left, congruent right, incongruent right) and this was subtracted from the RT for that trial type during the TMS experiment, on a trial-by-trial basis. This gave us the *difference* in RT for a specific trial type, when the participant was undergoing TMS. Therefore, all analyses of RTs looked at how the predictor variables changed the impact of TMS on RTs. Inspection of initial linear mixed-effects models showed that residuals were not normally distributed, caused by a rightward skew typical in reaction time data (Baayen & Milin, 2010; Ratcliff, 1993). Following Baayen & Milin (2010), data trimming and transformations, followed by model inspection, were explored to best normalize residuals for the models. The best results were achieved with a single cut-off value, excluding all reaction time differences longer than 400ms (the minimum reaction time difference was −389ms, so an upper cut-off was selected to mirror the minimum value). This removed 2.9% of the data.

To allow analysis of Scalp Location as a within subjects variable, we coded homologous scalp locations across left and right hemispheres together to look at the effect of scalp location on RT. For example, FP1 (left) and FP2 (right) were coded together, the two ATL sites were coded together, and so on.

There are three analyses. (1) A descriptive analysis of ratings across scalp locations and orientations. (2) Predicting RT by scalp location and orientation and (3) Predicting RT differences from subjective ratings. The second and third analyses were completed using linear mixed-effect models implemented in R (R Core Team, 2014); using the packages lme4 (Bates, Maechler, Bolker, & Walker, 2014), multcomp (Hothorn, Bretz, & Westfall, 2008), lmerTest (Kuznetsova, Brockhoff, & Bojesen, 2014), ggplot2 (Wickham, 2009) and corrplot (Wei, 2013). For these models, the reference level for scalp location was set to Cz/vertex, illustrating the effect of TMS across scalp locations relative to a commonly used control.

## 3.0 Results

### 3.1 Task performance

We compared error rates between Flanker and CRT tasks and found no significant difference (median errors Flanker=5 (range 0-44), median errors CRT=5.5 (range 0-66); Kruskal-Wallis=0.0230, df=1, p-value=0.879). Within task (Wilcoxon signed ranks test), there was no significant difference between congruent and incongruent conditions for the Flanker task (median errors congruent=1.5 (range 0-6), median errors incongruent=2 (range 0-41); W=35, p=0.264) and no significant difference for the CRT task (median errors congruent=3 (range 0-29), median errors incongruent=2.5 (range 0-37); W=49.5, p>0.900).

### 3.2 Scalp mapping of annoyance ratings and twitches

For participant ratings of subjective annoyance, pain, and twitches, and for the experimenter assessments of visible twitches, descriptive and inferential statistics were derived for each location, orientation, and hemisphere. A full account of these rich data is not possible here. Instead, we report the two dependent variables that we suggest are the most useful: The median ratings of muscle twitches and the mean effect of TMS on RT. Ratings of annoyance, pain, and twitches were strongly co-linear; twitches were most representative (see below). The effect of TMS on RT is likely the most useful measure for behavioural studies. The raw data are available at http://www.tms-smart.info for viewing and download, along with interactive maps of all available dependent variables and statistics, and a facility to find suggested control sites for each TMS location (10:20 system) and for individual MNI coordinates.

Median ratings of twitches (Figure 2, upper panels) varied from 0 (at superior sites on or adjacent to the midline) to 7 out of 10 (inferior frontal and anterior temporal sites). Ratings were symmetrical across the midline, and included four peaks – over the left and right anterior temporal/inferior frontal cortices, and over the left and right lateral occipital cortices.

**Figure 2:**
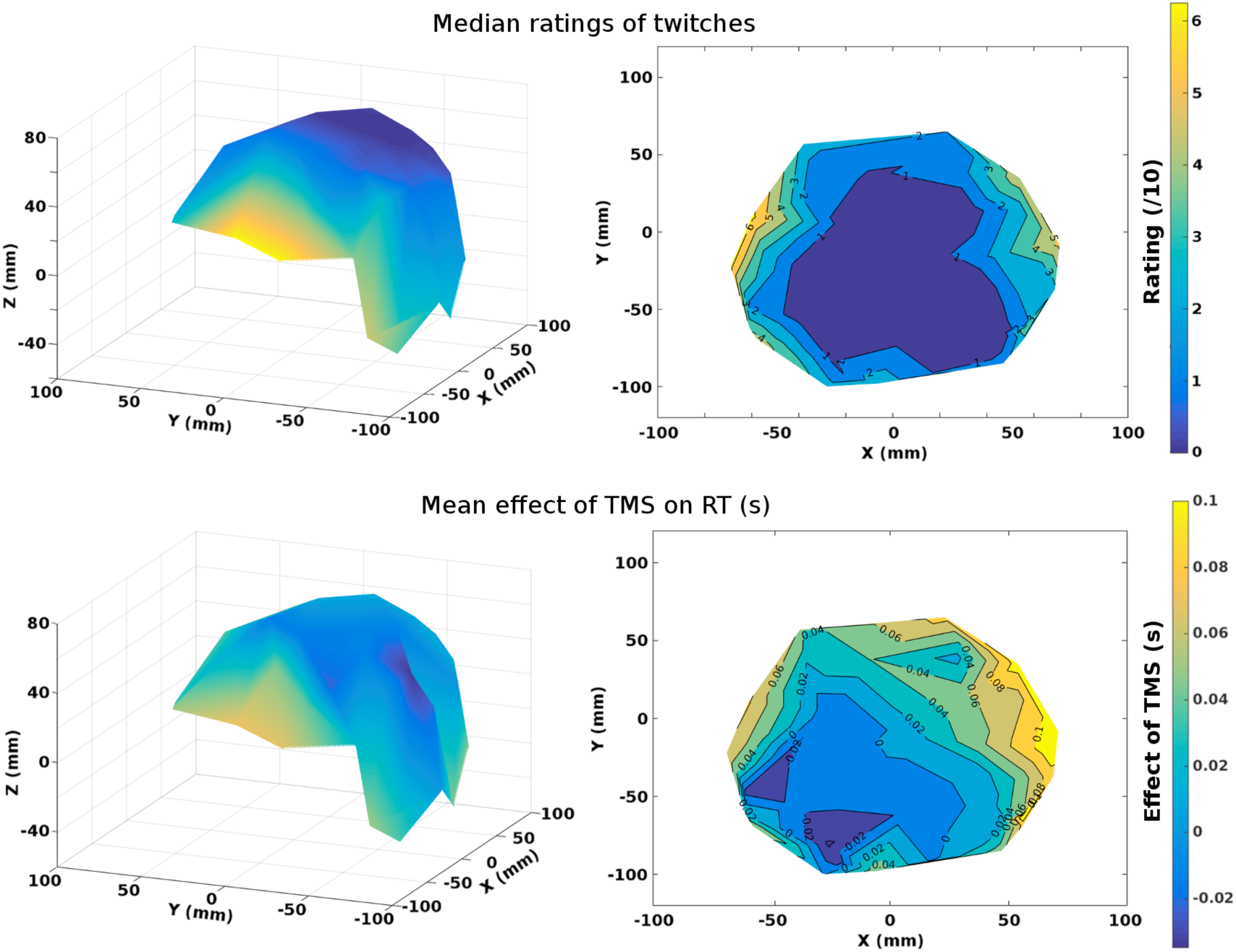
Interpolated maps of the effects of TMS location on the median participant rating of muscle twitches (upper panels) and the mean reaction time (RT, lower panels), averaged across the four coil orientations. Legend/Key: The maps in the left column show a 3D interpolation and rendering, viewed from behind the participant’s left ear, of the raw data mapped to the approximate Montreal Neurological Institute (MNI) coordinates (X, Y, Z, mm), as calculated in FSL, using a 12 degree-of-freedom affine transformation in FLIRT. The maps in the right column show a 2D plan view (from above) of the same data, and include the numerical values in the contour maps to show the scale. Dark blue shows low ratings (upper panels) or RT decrease (lower panels); while bright yellow shows higher ratings (upper panels) or RT increases (lower panels).

Mean effects of TMS on RT (Figure 2, lower panels) showed a broadly comparable distribution across the scalp, with the largest impairments in RT (up to 181ms) seen at inferior frontal, anterior temporal, and lateral occipital sites. The smallest changes in RT occurred along and adjacent to the superior midline, while TMS-induced improvements in RT (up to 61ms) were seen at superior parietal and occipital sites in the left hemisphere. The following section assesses the relationships between participant ratings and the effects of TMS on behaviour.

### 3.3 Predicting reaction times by scalp location and coil orientation

Random intercepts were fit for each participant to account for participant-by-participant variation in RTs (variance=1055ms). Following Barr et al (2013) we fit random effects to control for the within subjects variation in the effects of Scalp Location (variance=1360ms), Trial (trial within each scalp location, a number between 1-20; variance=425ms), Coil Orientation (variance=15ms), and Congruence (variance=581ms). To check that Scalp Location could be coded as homologous locations across Hemispheres, we tested the interaction between Scalp Location and Hemisphere. The interaction did not significantly improve model fit when compared against a model with Scalp Location as a main effect only (X^2^ goodness of fit=21.884, df=24, p=0.59). However, three scalp locations showed significant differences between hemispheres. CP5&6 (estimate=57.23ms, Standard Error=24.88ms, df=73, t=2.301, p=0.024), P7&8 (estimate=50.27ms, SE=25.00ms, df=75, t=2.011, p=0.048) and FC1&2 (estimate=55.54ms, SE=24.88, df=73, p=0.029). In all cases, RT costs under TMS were greater for the right hemisphere. Therefore, the interaction of Scalp Location and Hemisphere was retained in the final model as a control. To control for the effect of preceding trials (i.e. if the previous trial had been particularly painful, or an error trial) we included the reaction time from the previous trial. This was taken from within each block of 5 trials, so trial 1 had a value of 0 for the previous trial RT, trial 2 had the RT from trial 1, and so on, up to trial 5. This significantly improved model fit (X^2^=158.87, df=1, p<0.001; estimate=0.09, SE=0.01, df=861.7, t=12.57, p<0.001). The main effect of Task improved model fit (X^2^=4.443, df=1, p=0.035), the Flanker task showed a marginally greater impact of TMS on RTs than the CRT task (estimate=34.97ms, SE=16.99, df=17, t=2.06, p=0.055). The main effect of Congruence was not significant (p=0.366) nor did it improve model fit (X^2^=0.8785, df=1, p=0.35). There was no significant interaction between Task and Congruence (p>0.1) and the interaction term did not improve model fit (X^2^=1.92, df=2, p=0.38). The effect of Coil Orientation was not significant and did not improve model fit (X^2^=4.86, df=3, p=0.18); there were no significant differences between any TMS coil orientations (Tukey Contrasts all p>0.6). Coil Orientation and Congruence were not included as fixed effects predictors in the final model. With random effects noted as re(), the final model that best explained the data was:

RT Difference under TMS ∼ Scalp Location + Scalp Location x Hemisphere + Task + Previous RT + re(Participant) + re(Location x Participant) + re(Axes x Participant) + re(Congruence x Participant) + re(Trial x Participant)

The fixed effects alone had an R^2^ of 0.14, and the full model had an R^2^ of 0.43 (effect size of 0.80; Champely, 2016; Cohen et al., 2003; Selya et al., 2012; Lefcheck, 2015).

The effect of TMS on RTs varied across Scalp Locations, Figure 3 plots the mean RT difference data for each location, and Table 1 presents the estimated mean RT effect for each location (least squares means and standard errors, estimated difference against zero) and the approximate MNI location. Multiple comparisons (Tukey contrasts) comparing each Scalp Location to each other Scalp Location are provided in Appendix B. Here we present the estimated differences between each location and Cz/vertex, as this is a commonly used control location in TMS studies. Researchers wishing to re-reference the data to other sites and/or orientations can do so with the freely-available raw dataset at http://www.tms-smart.info. A number of sites did not differ significantly from Cz/vertex, these were C3&4, CP1&2, CP5&6, AF, PZ, P3&4, P7&8, FZ, F3&4, FC1&2, OZ, 01&2, TPJ, Lateral Occipital 3&4, Lateral Inion, and Inion. Six sites showed a greater cost of TMS than Cz/vertex. These were FC5&6 (est. mean=42.62ms, SE=18.58ms, t(383)=2.293, p=0.02); Lateral Occipital 1&2 (est. mean=53.43ms, SE=18.66ms, t(388)=2.864, p=0.004); ATL (est. mean=7.01ms, SE=18.56ms, t(381)=3.781, p=0.0002); T7&8 (est. mean=39.55ms, SE=18.62ms, t(386)=2.124, p=0.034), FP (est. mean=43.47ms, SE=18.70ms, t(392)=2.325, p=0.02), and F7&8 (est. mean=59.69ms, SE=18.64ms, t(387)=3.203, p=0.001). One site had faster RTs under TMS, significantly faster when compared against Cz/vertex. This was PO3&4 (est. mean=-36.69ms, SE=18.56ms, t(380)=-1.977, p=0.048). Figure 4 presents the RT difference in milliseconds for each scalp location (TMS RT – No TMS RT).

**Figure 3:**
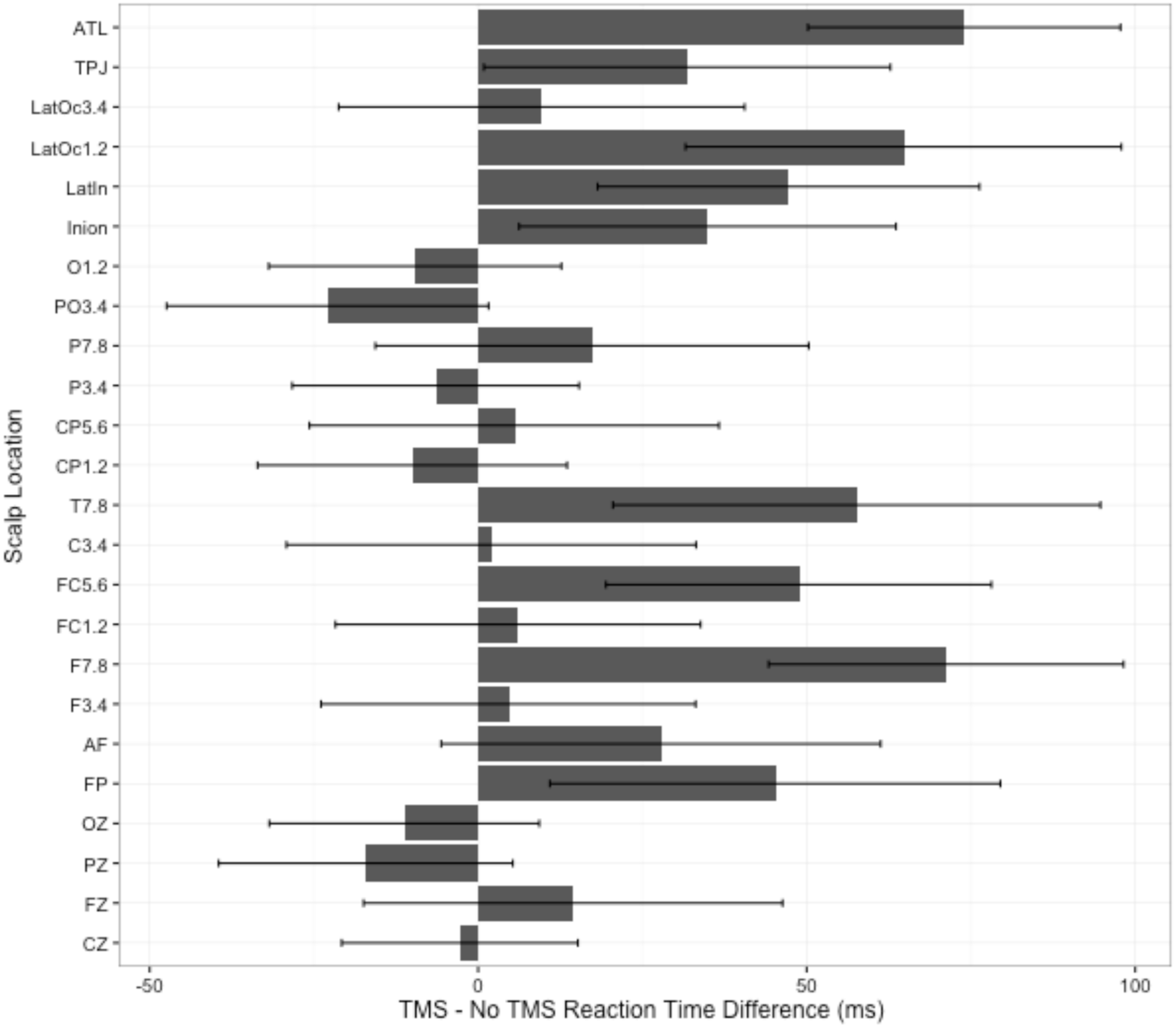
Effect of TMS by Scalp Location. Legend/Key: A positive difference means slower RTs under TMS, a negative difference means faster RTs under TMS. Error bars are 95% confidence intervals.

**Figure 4:**
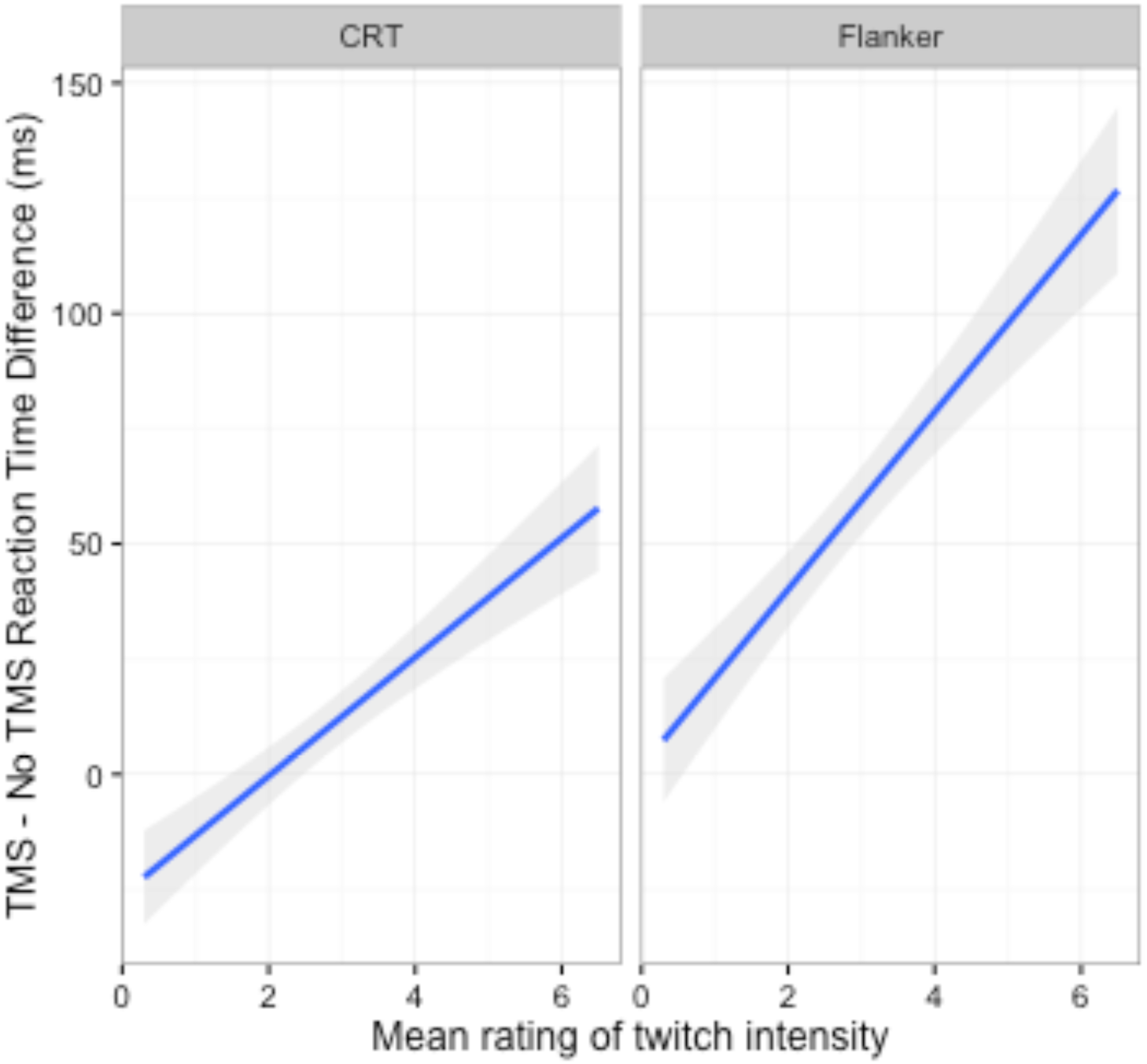
Reaction time difference (TMS – No TMS) by subjective rating of Twitch intensity for CRT and Flanker Tasks. Legend/Key: A positive value on the y axis means slower RTs under TMS. Error bars are 95% confidence intervals.

Table 1 presents least squares means for the effect of TMS on RTs, comparing each site against zero. The sites showing the biggest costs were frontal and lateral sites, some of which showed very high RT costs for TMS. The sites with the largest slowing of RTs under TMS were the ATL (∼75ms cost), F7&8 (∼70ms cost), Lateral-Occipital 1&2 (∼65ms cost) and T7&8 (∼55ms cost). The sites with the smallest cost were sites in the midline, some of which showed negligible changes. The standard error across sites was around 10ms, and the following sites showed a less than 10ms slowing under TMS: Cz (∼2ms), F3&4 (∼4.5ms), FC1&2 (∼6ms), C3&4 (∼3ms), CP5&6 (∼5ms). The complete least squares means broken down by hemisphere and location are provided in Appendix B.

### 3.4 Predicting reaction times by ratings of annoyance, pain, twitches, and by visible twitches

To test whether Twitch Ratings are a better explanation for the variation in RTs across Scalp Locations, we added Twitch Ratings as a fixed effect predictor to the existing model (detailed above). This significantly improved model fit (X^2^(1)= 28.81, p<0.001) and showed a significant main effect of Twitch Ratings (est. mean=14.99ms, SE=2.77, df=5411, t=5.406, p<0.0001). For every increment in rated Twitch intensity, there was an RT cost of around 15ms under TMS. We tested the interaction between Twitch Ratings and Task, to explore whether subjective discomfort differentially affected the two tasks. The interaction was significant (est. mean=7.75ms, SE=2.26ms, df=557, t=3.43, p=0.0007), with a greater cost in RTs for the Flanker task than the CRT task. The interaction term significantly improved model fit (X^2^(1)=12.454, p<0.0005). To illustrate the interaction, Figure 4 plots the effect of Twitch Ratings on reaction times (ms), by task.

Once Twitch Rating was included in the model, significant variation in RTs as predicted by Scalp Location was substantially reduced. In this model, only one Scalp Location showed a significant difference to Cz. This was P03&4 (est. mean=-39.77, SE=18.49, df=381, t=-2.150, p<0.05), which retained significantly speeded RTs under TMS relative to Cz. All Scalp Locations which had shown slowing under TMS relative to CZ in the previous model were no longer significant. For this model, the fixed effects had an R^2^ of 0.14 (effect size 0.15) and the full model (i.e., both fixed and random effects) had an R^2^ of 0.43 (effect size of 0.81).

Appendix D provides details of a parallel analysis that uses scalp to brain distance instead of Scalp Location, where we found identical results. Thus, when subjective discomfort is taken into account (i.e., the inclusion of Twitch Ratings as a predictor), neither Scalp Location nor scalp to brain distance predict the cost of TMS on RTs.

To check whether individual variation in resting motor threshold (RMT) influenced the results, we added each participant’s RMT as a predictor to this model, both as a main effect (i.e. does a higher or lower RMT change how much TMS influences overall RTs in the task) and with an interaction between RMT and Scalp Location (i.e. does the influence of TMS on RTs change for particular scalp locations when the participant has a higher or lower RMT?). It could be argued that individuals with lower RMTs would be more likely to experience cortical stimulation from TMS (fixed at 50% of maximum stimulator output). We might see that a lower RMT would lead to greater changes in RTs under TMS (a main effect) or particular cortical locations would show greater RT changes under TMS for individuals with lower RMT (an interaction). We found no such effects, that is, there was no significant main effect of RMT and no interaction between RMT and Scalp Location.

## 4.0 Discussion

When mapped across the scalp, subjective ratings and differences in RTs showed minimal changes along the midline but high interference from TMS at inferior frontal, anterior temporal, and lateral occipital sites. This confirms previous reports that TMS is more uncomfortable – and often painful – over frontal scalp locations (Abler et al., 2005; Loo et al., 2008; Machii et al., 2006; Maizey et al., 2013; Wassermann, 1998). This experiment is the first to provide data that systematically maps these effects across the scalp.

We found significant differences in how TMS affected reaction times when the pulse was applied to different scalp locations. Analyses compared each scalp location against Cz/vertex, a commonly used control site in TMS studies (e.g., Jung et al., 2016), and six locations produced longer RTs under TMS as compared to Cz/vertex. These results are in line with the argument that TMS over Cz seemingly does not affect behaviour (e.g., Jung et al., 2016; Silvanto et al., 2008), as Cz showed no substantial change in RTs when on-line TMS was applied. This makes Cz/Vertex a suitable control for other similarly benign scalp locations (e.g., FZ, FC1&2, C3&4, etc., see Table 1 in Results). However Cz/Vertex will only control for limited dimensions of TMS (e.g., the audible click, presence of the experimenter) when compared against scalp locations that are more uncomfortable under TMS (e.g., Lateral-Occipital, ATL, T7&8).

We found that subjective ratings of annoyance, pain, and twitches were highly correlated; this is perhaps unsurprising given that participants provided these ratings consecutively. It does however support previous findings that the discomfort caused by TMS is primarily due to the effect of the TMS pulse on the skin, nerves, and muscle of the scalp (Abler et al., 2005; Arana et al., 2008; Loo et al., 2000). Once ratings of muscle twitches was added as a predictor variable, the effect of Scalp Location disappeared for all areas where TMS caused longer RTs. This means that the increase in RTs seen for many Scalp Locations under TMS is due to the fact that these locations are more uncomfortable under TMS. This finding did not change when we included scalp-to-cortex distance rather than scalp location. Abler and colleagues (2005) found a positive correlation between the number of errors on a task and subjective ratings of discomfort caused by TMS. Our data provides further support for the relationship between discomfort under TMS and changes in behavioural performance.

One question that we have been asked is “How do you know it wasn’t brain stimulation that caused the differences in behavioural response times?” It is likely that cortical stimulation did occur at most TMS sites. However, the degree of cortical stimulation across different locations was, at best, poorly controlled due to varying scalp to brain distances (Stokes et al, 2005; Sack et al, 2009). When compared against other means of localising cortical sites for stimulation (i.e. fMRI guided, MRI guided and Talairach) the 10:20 anatomical system shows the smallest effect size for effects of TMS (versus sham) on RTs (Sack et al., 2009). We found no effect of scalp to brain distance and no effect of resting motor threshold on RT once subjective discomfort was accounted for. The most parsimonious explanation for the cause of the clear and consistent effects of TMS on RTs is peripheral stimulation and discomfort.

The one scalp location that showed speeded RTs under TMS (PO3&4) remained significant in the analysis that included subjective ratings. Previous studies have shown that TMS over parietal sites affects visual attention (Chambers & Heinen, 2010; Taylor & Thut, 2012), so this result could be explained by cortical stimulation broadly enhancing visual attention or orienting during the task. Interestingly, for the parietal sites with faster RTs under TMS (PO3&4) median ratings of annoyance, pain, and twitches were close to zero. Therefore, at these sites the TMS pulse at stimulus onset may have acted as a general alerting signal that reduced RTs (Drager et al., 2004; Duecker et al., 2013; Duecker & Sack, 2013; Marzi et al., 1998; Nikouline et al., 1999; Sawaki et al., 1999). Our data is not able to distinguish between these two explanations.

The data also showed that greater discomfort (i.e., subjective ratings) produces a greater cost on task performance, and this cost differed by task. For every unit increase in annoyance there was a 14ms increase in RTs for the CRT task, and a 21ms increase in RTs for the Eriksen flanker task. Thus, more difficult tasks are affected more by the TMS-induced discomfort. Models explained ∼45% of the variance in RTs. This is comparable, for example, to the size of the effect of scalp to brain distance on resting motor threshold (Stokes et al., 2007). Because of this relationship control tasks need to be carefully matched to experimental tasks in TMS experiments in order for confounds to be managed (Rossi et al., 2009). We should stress that our findings apply only to online single pulse TMS studies (i.e., the conditions that we tested), where the TMS pulse is delivered during task performance, close to stimulus presentation and the preparation and execution of a behavioural response. Whilst online and offline rTMS, paired-pulse, quadripulse, and other TMS protocols are all likely to induce similar scalp sensations, their influence on task performance needs to be determined.

We did not measure the participant’s ratings of auditory or visual interference of TMS at each site (although participants’ wore ear-plugs to mitigate the auditory interference). For sites close to the ears, the TMS-related sounds would have been louder and more lateralized than sites further away. For sites close to the eyes, participants’ peripheral vision would have been partly occluded by the TMS coil. These variables may have explained additional variance in the data, and should be studied further.

The data we have presented systematically maps the peripheral effects of single-pulse TMS across the scalp. We cannot generalise these results to other TMS stimulation parameters, but note that studies which use bursts of TMS will tend to result in higher levels of subjective discomfort, which can lead to even more “side-effects” and annoyance. As noted above, best practice in TMS studies is to have more than one control condition for a reliable, specific effect of TMS (Sandrini et al., 2011). Our data provides a tool to select control sites for on-line TMS experiments (http://www.tms-smart.info) when researchers want to control for annoyance, pain and twitches. In an ideal case, the control site will be a different brain region/scalp location, selected to match for subjective ratings (annoyance, pain, or twitches), or the effect on RTs. For example, if the experimental TMS site is over the right TPJ with the coil handle oriented approximately 45 degrees behind the inter-aural axis (SE-NW, median rating of twitches=3), suitable control sites using the same coil orientation, include the right lateral inion (7.91cm away, median twitches=3), the right lateral occipital 2 (6.81cm away, median twitches=3), or FC6 (6.80cm away, median twitches=2.5). The tools on the website allow you to customise the search for control locations using all our available data, the 10:20 system, and/or MNI co-ordinates.

Once target and control sites have been selected, we would then recommend a systematic pilot exploration of the effects of TMS intensity and coil orientation on participant-reported levels of annoyance, twitches, and pain, and on RT. We have provided an example of a systematic exploration that could be used to select appropriate TMS intensities for different sites (Appendix C). Interestingly, for scalp locations shown to be the most annoying from our initial data, the effect of TMS intensity on ratings was linear (see Appendix C). Our data provides a tool to select control sites for on-line TMS experiments (http://www.tms-smart.info) when researchers want to control for annoyance, pain and twitches. The tools on the website allow you to customise the search for control locations using all our available data, the 10:20 system, and/or MNI co-ordinates. We hope this data and the online resource will be useful for future TMS studies to properly control for the peripheral effects of TMS, particularly when those peripheral effects are extremely annoying.

## Acknowledgements

We would like to thank Arran Reader, Nergis Yüksel and Ainara Jauregi for help with experiment preparation, data collection and literature searches. We would also like to thank Elisabeth Volke for help with data collation and preparation. We would like to thank previous reviewers of the manuscript for robust comments that have improved and extended our analyses. This project was funded by the Research Group Pump Priming scheme (Language & Cognition, Perception & Action) awarded to the authors from the University of Reading School of Psychology and Clinical Language Sciences.

## Appendix A

**Table A1:** Task by participant summary of mean RT, proportion correct and congruency effects.

| Task | Participant | Mean correct RT (ms) | Proportion correct | Congruency effect (Incongruent – Congruent, ms) |
| --- | --- | --- | --- | --- |
| Flanker | 1 | 437 | 0.99 | 14.6 |
|  | 2 | 428 | 0.91 | 49.9 |
|  | 3 | 563 | 0.99 | 44.7 |
|  | 4 | 644 | 0.99 | 42.1 |
|  | 5 | 578 | 0.99 | -6.2 |
|  | 6 | 467 | 0.97 | 37.0 |
|  | 7 | 558 | 1.00 | 13.5 |
|  | 8 | 592 | 0.96 | 44.5 |
|  | 9 | 622 | 1.00 | 40.2 |
|  | 10 | 604 | 0.97 | 49.4 |
| **Mean (SD)** |  | **549 (77.6)** | **0.98 (0.03)** | **33.0 (18.9)** |
| CRT | 11 | 412 | 0.96 | 12.1 |
|  | 12 | 330 | 1.00 | 3.8 |
|  | 13 | 295 | 0.86 | 3.7 |
|  | 14 | 417 | 0.98 | 18.4 |
|  | 15 | 385 | 0.98 | 5.5 |
|  | 16* | 629 | 0.99 | 20.8 |
|  | 17 | 286 | 0.98 | -3.3 |
|  | 18 | 494 | 0.99 | 26.4 |
|  | 19* | 431 | 0.99 | 23.9 |
|  | 20* | 485 | 0.99 | 50.5 |
| **Mean (SD)** |  | **416 (103)** | **0.97 (0.04)** | **16.2 (15.6)** |

\* Participants who responded to arrow direction rather than side of presentation, data were recoded to reflect their responses.

## Appendix B

### Multiple comparisons for all scalp locations

Adjusted P values using Tukey HSD to control for family wise error rate.

For each comparison, the null hypothesis is that the difference between the Test Location and Reference is 0.

A positive estimate means that the Test Location shows a greater cost of TMS (i.e. longer RTs under TMS) than the Reference. A negative estimate means that the Test Location shows less cost (i.e. shorter RTs under TMS) than the Reference.

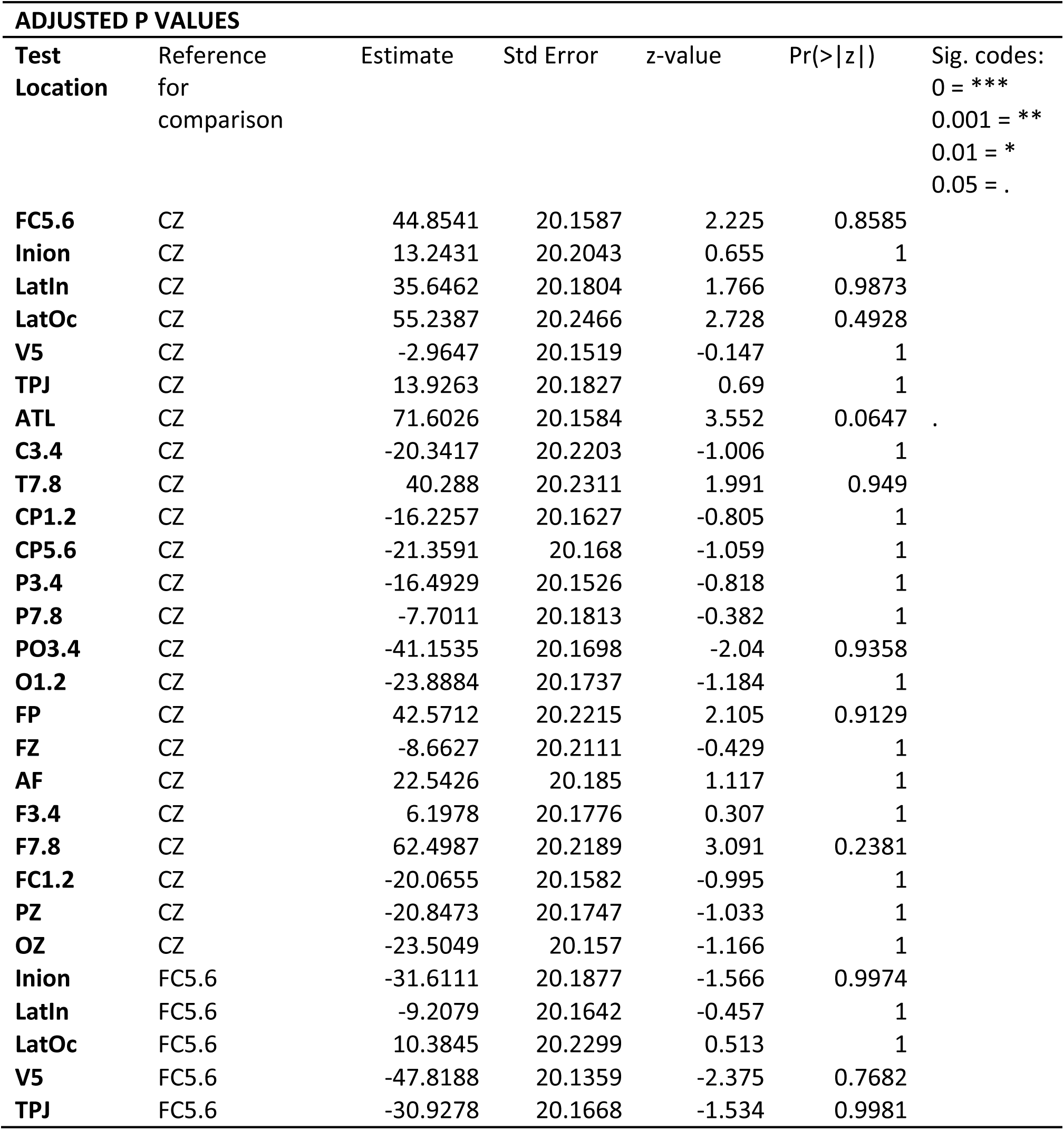
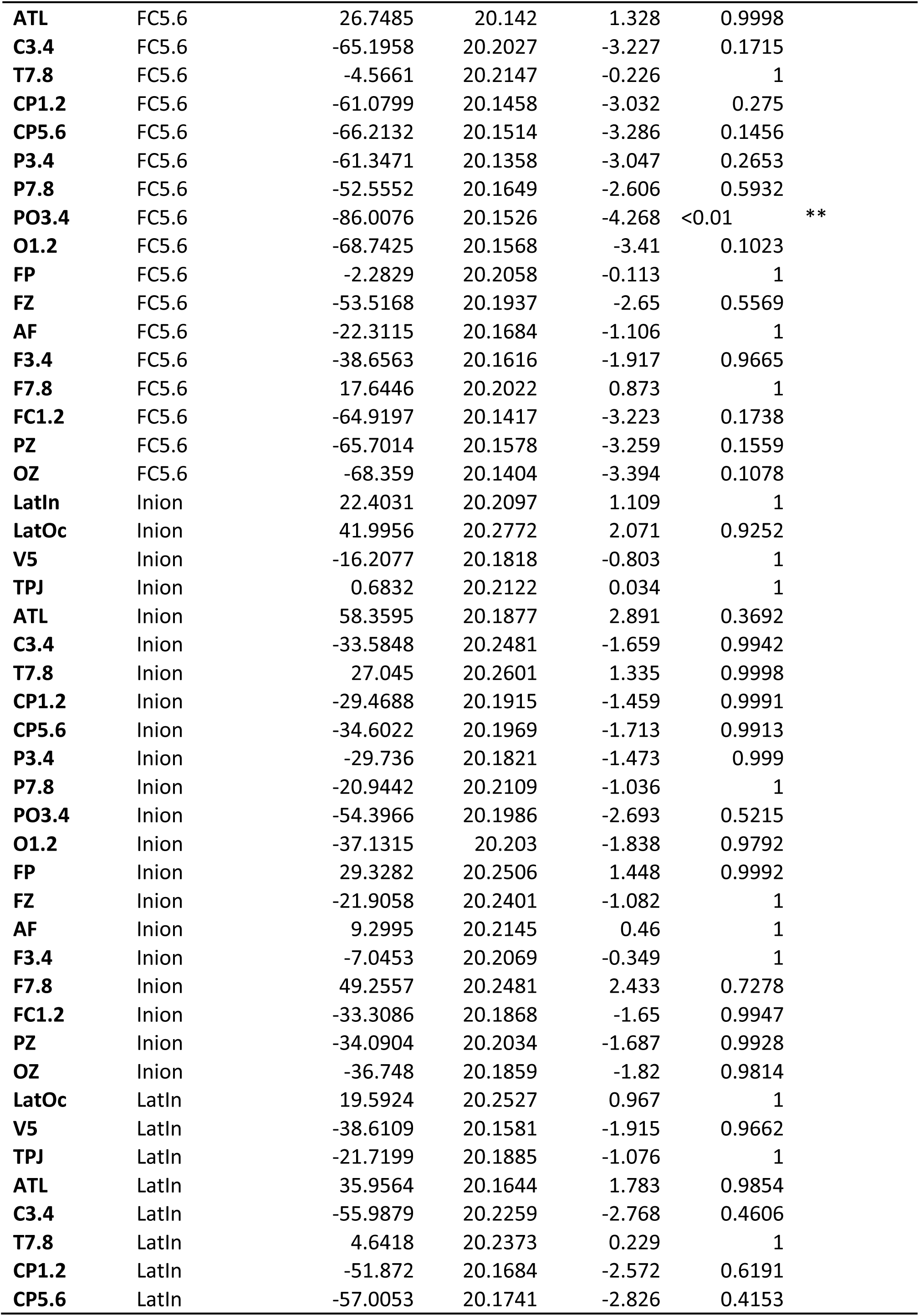
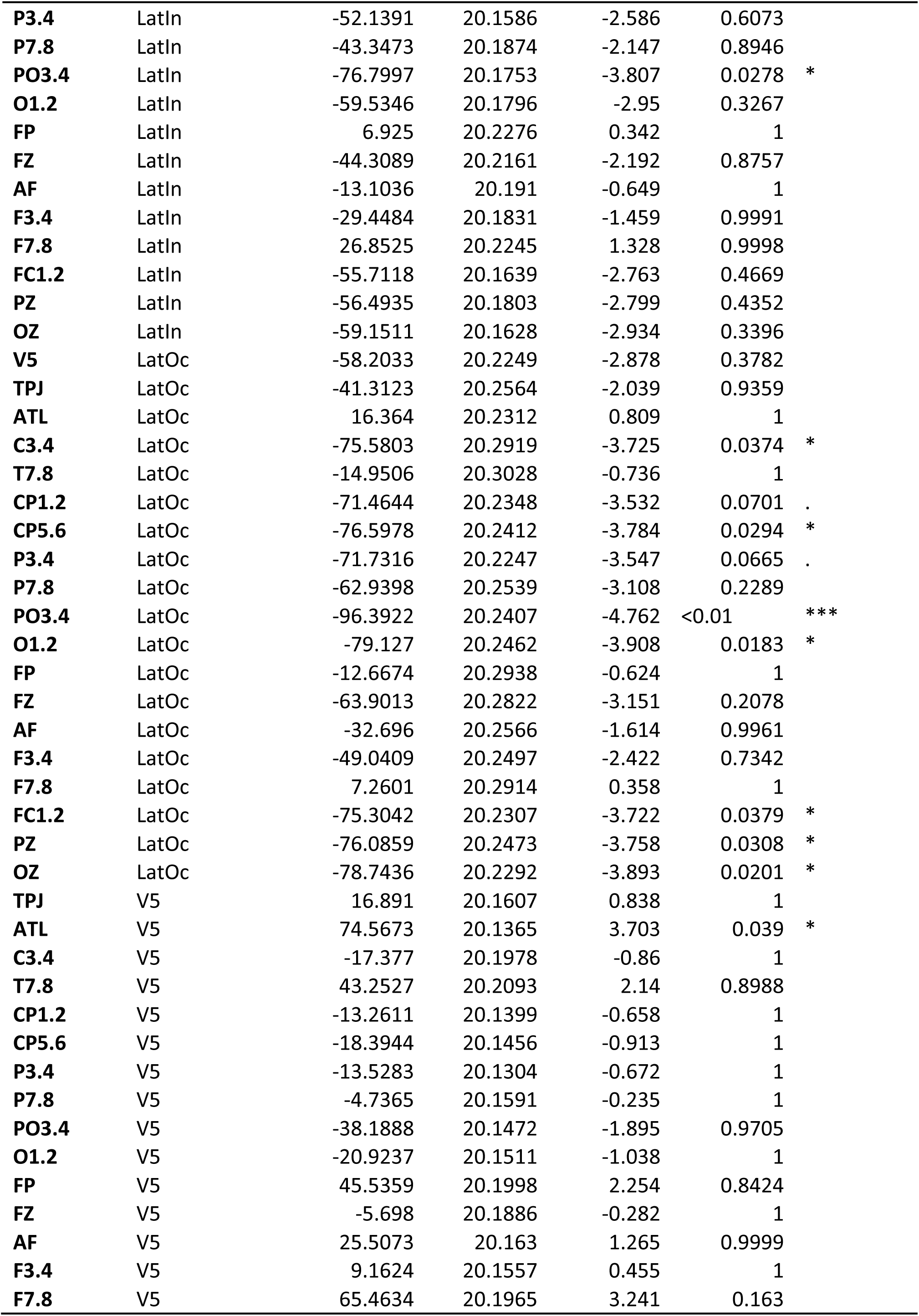
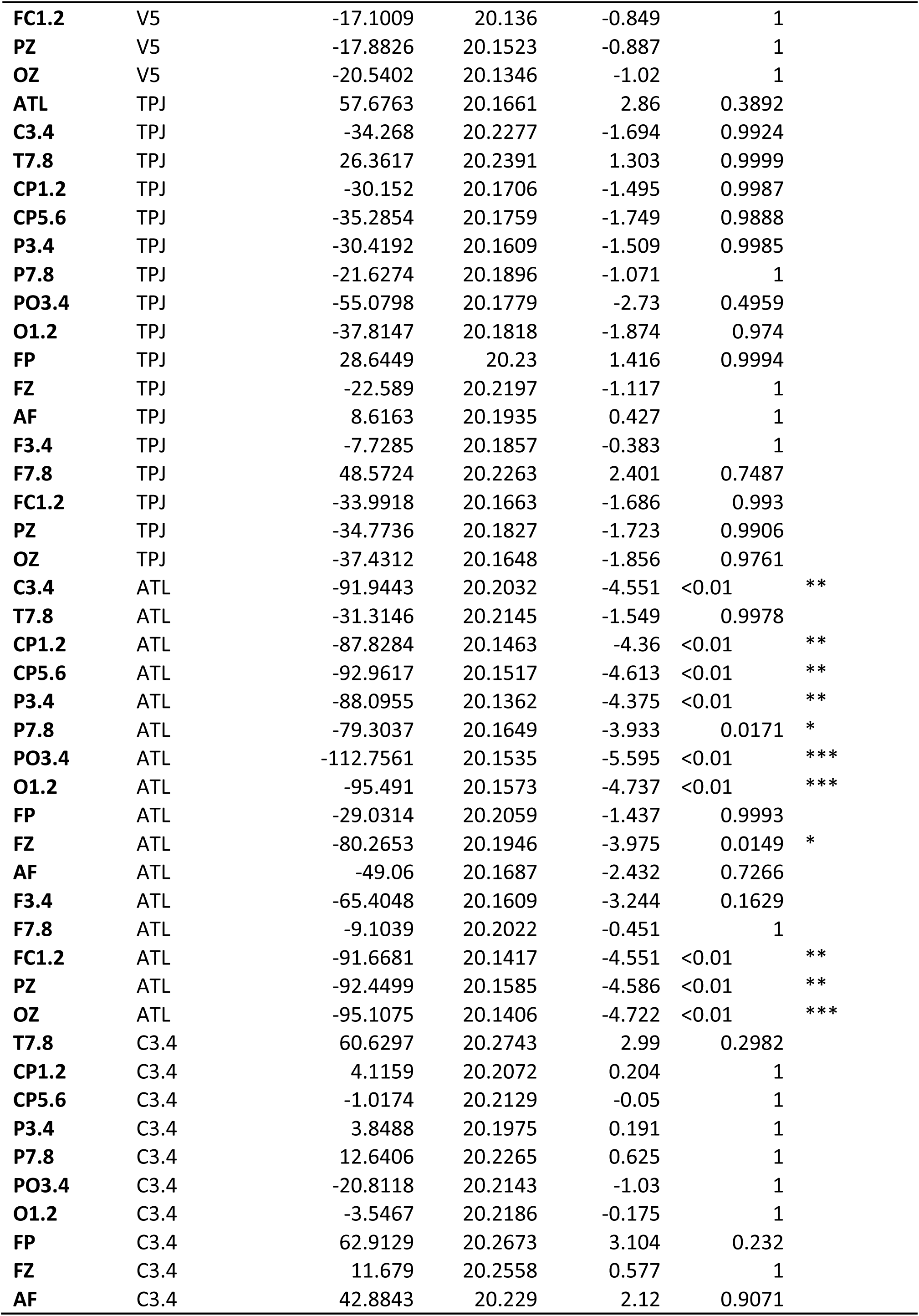
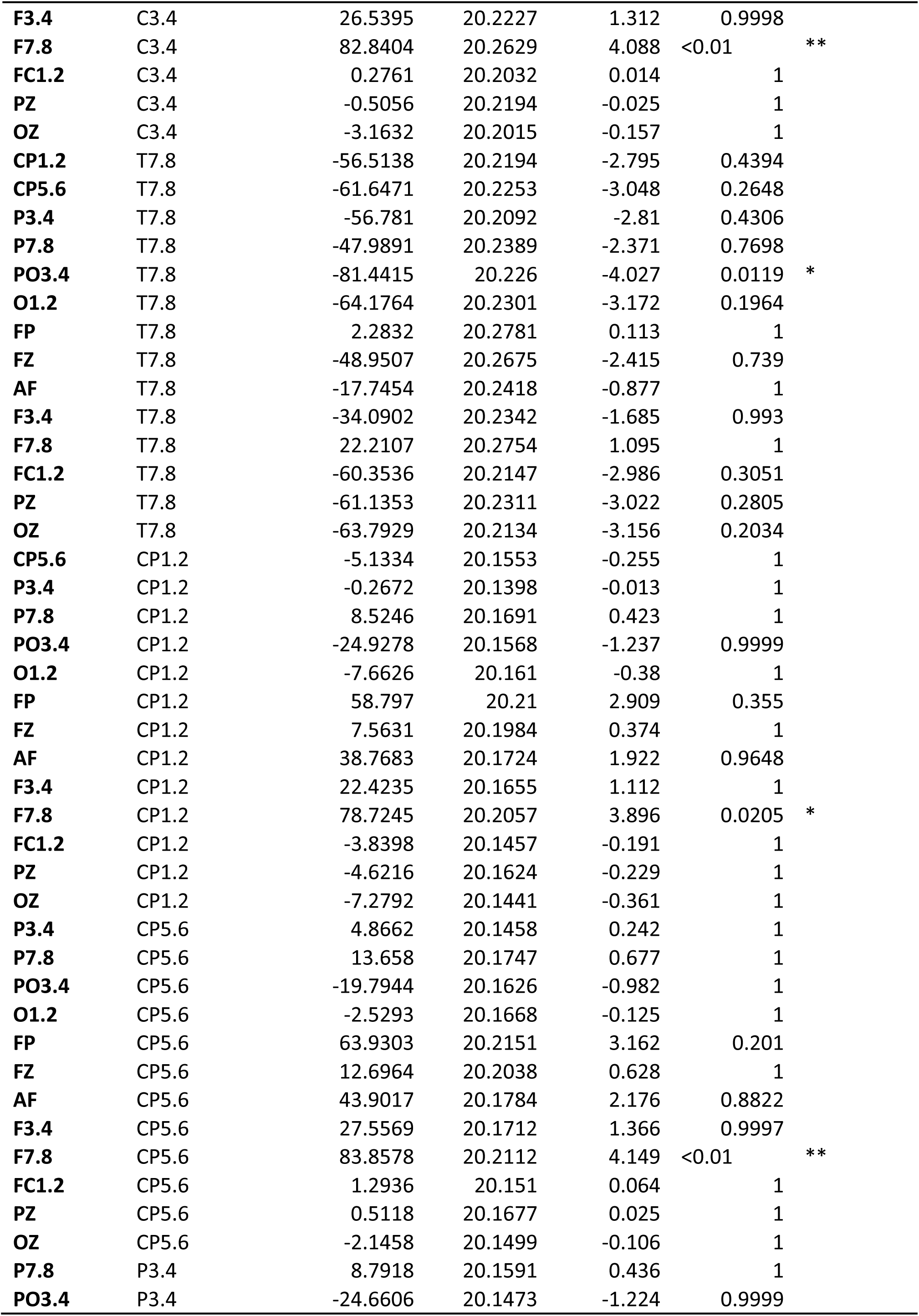
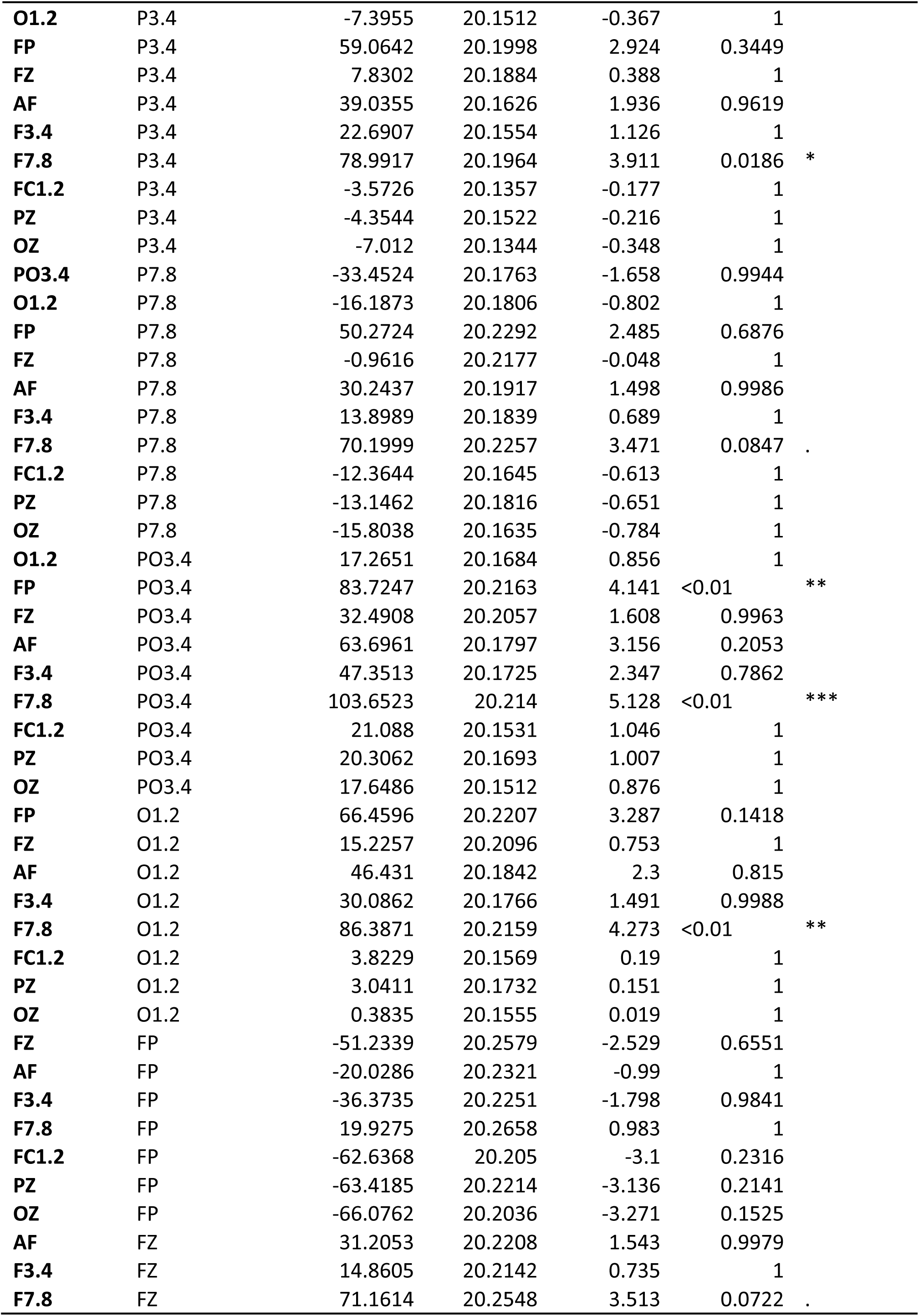
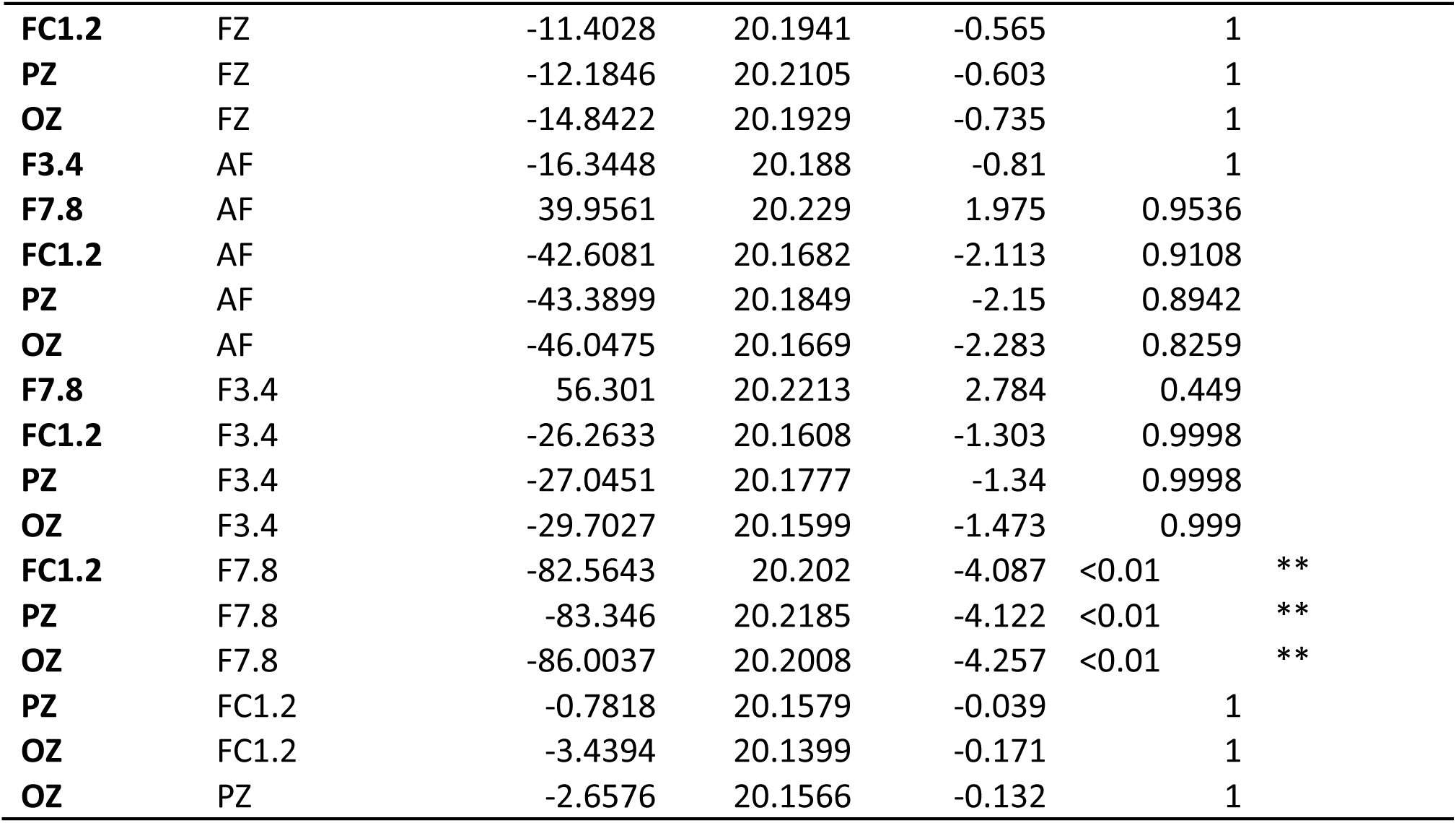

Appendix B Least squares means for all scalp locations and scalp locations x hemisphere.

Uncorrected p values and Bonferroni correction are reported.

For each comparison, the null hypothesis is that the value for each Scalp Location is 0.

A positive estimate means that the Scalp Location shows a greater cost of TMS (i.e. longer RTs under TMS). A negative estimate means that the Scalp Location shows an advantage under TMS (i.e. shorter RTs under TMS).

| Scalp Location | Estimate | St. Err | DF | t-value | Lower CI | Upper CI | Uncorrected p value | Sig. with Bonferroni corrected p < 0.002 |
| --- | --- | --- | --- | --- | --- | --- | --- | --- |
| **CZ** | -1.955 | 13.538 | 73.3 | -0.14 | -28.934 | 25.02 | 0.886 |  |
| **FC5.6** | 48.482 | 13.557 | 73.8 | 3.58 | 21.467 | 75.5 | 6.00E-04 | * |
| **Inion** | 35.381 | 13.553 | 73.7 | 2.61 | 8.374 | 62.39 | 0.011 |  |
| **LatIn** | 47.743 | 13.55 | 73.6 | 3.52 | 20.741 | 74.74 | 7.00E-04 | * |
| **LatOc** | 65.332 | 13.604 | 74.8 | 4.8 | 38.229 | 92.43 | 2E-15 | * |
| **V5** | 9.928 | 13.547 | 73.5 | 0.73 | -17.067 | 36.92 | 0.466 |  |
| **TPJ** | 31.986 | 13.585 | 74.4 | 2.35 | 4.918 | 59.05 | 0.021 |  |
| **ATL** | 74.564 | 13.552 | 73.7 | 5.5 | 47.559 | 101.57 | 2E-15 | * |
| **C3.4** | 2.917 | 13.581 | 74.3 | 0.21 | -24.143 | 29.98 | 0.831 |  |
| **T7.8** | 56.49 | 13.623 | 75.2 | 4.15 | 29.353 | 83.63 | 1.00E-04 | * |
| **CP1.2** | -10.46 | 13.536 | 73.3 | -0.77 | -37.435 | 16.52 | 0.442 |  |
| **CP5.6** | 5.44 | 13.548 | 73.6 | 0.4 | -21.557 | 32.44 | 0.689 |  |
| **P3.4** | -6.252 | 13.533 | 73.2 | -0.46 | -33.222 | 20.72 | 0.645 |  |
| **P7.8** | 16.892 | 13.552 | 73.6 | 1.25 | -10.112 | 43.9 | 0.216 |  |
| **PO3.4** | -22.888 | 13.532 | 73.2 | -1.69 | -49.857 | 4.08 | 0.095 |  |
| **O1.2** | -9.569 | 13.53 | 73.2 | -0.71 | -36.533 | 17.39 | 0.482 |  |
| **FP** | 45.827 | 13.603 | 74.8 | 3.37 | 18.727 | 72.93 | 0.001 | * |
| **FZ** | 13.96 | 13.578 | 74.2 | 1.03 | -13.094 | 41.01 | 0.307 |  |
| **AF** | 27.506 | 13.547 | 73.5 | 2.03 | 0.511 | 54.5 | 0.046 |  |
| **F3.4** | 4.604 | 13.553 | 73.7 | 0.34 | -22.403 | 31.61 | 0.735 |  |
| **F7.8** | 71.272 | 13.562 | 73.9 | 5.26 | 44.248 | 98.3 | 2E-15 | * |
| **FC1.2** | 6.28 | 13.542 | 73.4 | 0.46 | -20.707 | 33.27 | 0.644 |  |
| **PZ** | -16.83 | 13.533 | 73.2 | -1.24 | -43.799 | 10.14 | 0.218 |  |
| **OZ** | -11.081 | 13.531 | 73.2 | -0.82 | -38.047 | 15.89 | 0.415 |  |

| Scalp Location x Hemisphere | Hemisphere | Estimate | St. Err | DF | t-value | Lower CI | Upper CI | Uncorrected p value | Sig. with Bonferroni corrected p < 0.001 |
| --- | --- | --- | --- | --- | --- | --- | --- | --- | --- |
| **CZ** | Left | -4.112 | 19.149 | 73.4 | -0.21 | -42.273 | 34.05 | 0.831 |  |
| **FC5.6** | Left | 40.742 | 19.132 | 73.1 | 2.13 | 2.614 | 78.87 | 0.037 |  |
| **Inion** | Left | 9.131 | 19.18 | 73.9 | 0.48 | -29.087 | 47.35 | 0.635 |  |
| **LatIn** | Left | 31.535 | 19.156 | 73.5 | 1.65 | -6.638 | 69.71 | 0.104 |  |
| **LatOc** | Left | 51.127 | 19.226 | 74.6 | 2.66 | 12.824 | 89.43 | 0.01 |  |
| **V5** | Left | -7.076 | 19.126 | 73 | -0.37 | -45.194 | 31.04 | 0.713 |  |
| **TPJ** | Left | 9.815 | 19.158 | 73.5 | 0.51 | -28.362 | 47.99 | 0.61 |  |
| **ATL** | Left | 67.491 | 19.132 | 73.1 | 3.53 | 29.362 | 105.62 | 7.00E-04 | * |
| **C3.4** | Left | -24.453 | 19.197 | 74.1 | -1.27 | -62.702 | 13.8 | 0.207 |  |
| **T7.8** | Left | 36.176 | 19.209 | 74.3 | 1.88 | -2.096 | 74.45 | 0.064 |  |
| **CP1.2** | Left | -20.337 | 19.136 | 73.2 | -1.06 | -58.474 | 17.8 | 0.291 |  |
| **CP5.6** | Left | -25.471 | 19.142 | 73.3 | -1.33 | -63.618 | 12.68 | 0.187 |  |
| **P3.4** | Left | -20.605 | 19.126 | 73 | -1.08 | -58.722 | 17.51 | 0.285 |  |
| **P7.8** | Left | -11.813 | 19.156 | 73.5 | -0.62 | -49.987 | 26.36 | 0.539 |  |
| **PO3.4** | Left | -45.265 | 19.144 | 73.3 | -2.36 | -83.416 | -7.11 | 0.021 |  |
| **O1.2** | Left | -28 | 19.148 | 73.4 | -1.46 | -66.159 | 10.16 | 0.148 |  |
| **FP** | Left | 38.46 | 19.199 | 74.2 | 2 | 0.206 | 76.71 | 0.049 |  |
| **FZ** | Left | -12.774 | 19.187 | 74 | -0.67 | -51.006 | 25.46 | 0.508 |  |
| **AF** | Left | 18.431 | 19.16 | 73.6 | 0.96 | -19.75 | 56.61 | 0.339 |  |
| **F3.4** | Left | 2.086 | 19.152 | 73.5 | 0.11 | -36.081 | 40.25 | 0.914 |  |
| **F7.8** | Left | 58.387 | 19.196 | 74.1 | 3.04 | 20.14 | 96.63 | 0.003 |  |
| **FC1.2** | Left | -24.177 | 19.132 | 73.1 | -1.26 | -62.306 | 13.95 | 0.21 |  |
| **PZ** | Left | -24.959 | 19.149 | 73.4 | -1.3 | -63.12 | 13.2 | 0.197 |  |
| **OZ** | Left | -27.617 | 19.13 | 73.1 | -1.44 | -65.742 | 10.51 | 0.153 |  |
| **CZ** | Right | 0.202 | 19.142 | 73.3 | 0.01 | -37.944 | 38.35 | 0.992 |  |
| **FC5.6** | Right | 56.221 | 19.214 | 74.4 | 2.93 | 17.94 | 94.5 | 0.004 |  |
| **Inion** | Right | 61.63 | 19.154 | 73.5 | 3.22 | 23.461 | 99.8 | 0.002 |  |
| **LatIn** | Right | 63.951 | 19.17 | 73.7 | 3.34 | 25.752 | 102.15 | 0.001 | * |
| **LatOc** | Right | 79.537 | 19.253 | 75 | 4.13 | 41.182 | 117.89 | 1.00E-04 | * |
| **V5** | Right | 26.933 | 19.191 | 74 | 1.4 | -11.305 | 65.17 | 0.165 |  |
| **TPJ** | Right | 54.157 | 19.268 | 75.2 | 2.81 | 15.775 | 92.54 | 0.006 |  |
| **ATL** | Right | 81.638 | 19.2 | 74.2 | 4.25 | 43.383 | 119.89 | 1.00E-04 | * |
| **C3.4** | Right | 30.287 | 19.217 | 74.4 | 1.58 | -8 | 68.57 | 0.119 |  |
| **T7.8** | Right | 76.804 | 19.322 | 76 | 3.97 | 38.321 | 115.29 | 2.00E-04 | * |
| **CP1.2** | Right | -0.582 | 19.149 | 73.4 | -0.03 | -38.742 | 37.58 | 0.976 |  |
| **CP5.6** | Right | 36.351 | 19.177 | 73.8 | 1.9 | -1.86 | 74.56 | 0.062 |  |
| **P3.4** | Right | 8.1 | 19.151 | 73.4 | 0.42 | -30.065 | 46.26 | 0.674 |  |
| **P7.8** | Right | 45.597 | 19.174 | 73.8 | 2.38 | 7.391 | 83.8 | 0.02 |  |
| **PO3.4** | Right | -0.511 | 19.132 | 73.1 | -0.03 | -38.64 | 37.62 | 0.979 |  |
| **O1.2** | Right | 8.862 | 19.121 | 73 | 0.46 | -29.246 | 46.97 | 0.644 |  |
| **FP** | Right | 53.194 | 19.275 | 75.3 | 2.76 | 14.798 | 91.59 | 0.007 |  |
| **FZ** | Right | 40.694 | 19.218 | 74.4 | 2.12 | 2.406 | 78.98 | 0.038 |  |
| **AF** | Right | 36.582 | 19.156 | 73.5 | 1.91 | -1.592 | 74.76 | 0.06 |  |
| **F3.4** | Right | 7.121 | 19.18 | 73.9 | 0.37 | -31.098 | 45.34 | 0.712 |  |
| **F7.8** | Right | 84.157 | 19.164 | 73.6 | 4.39 | 45.969 | 122.35 | 2E-15 | * |
| **FC1.2** | Right | 36.737 | 19.172 | 73.8 | 1.92 | -1.466 | 74.94 | 0.059 |  |
| **PZ** | Right | -8.701 | 19.127 | 73.1 | -0.45 | -46.821 | 29.42 | 0.65 |  |
| **OZ** | Right | 5.455 | 19.142 | 73.3 | 0.28 | -32.691 | 43.6 | 0.776 |  |

## Appendix C / Supplementary materials 3: Effect of varying TMS intensity on ratings of annoyance, pain, and muscle twitches

Following reviewers’ comments on an earlier version of this manuscript, we studied, in one volunteer, the effect of varying TMS intensity on subjective ratings of annoyance, pain, and twitches, and on visible twitches. This approach could be used to further refine the choice of control site for an experiment.

### Methods

From the original dataset, we chose 6 scalp locations that spanned the full range of median twitch ratings (Table C1).

**Table C1:**
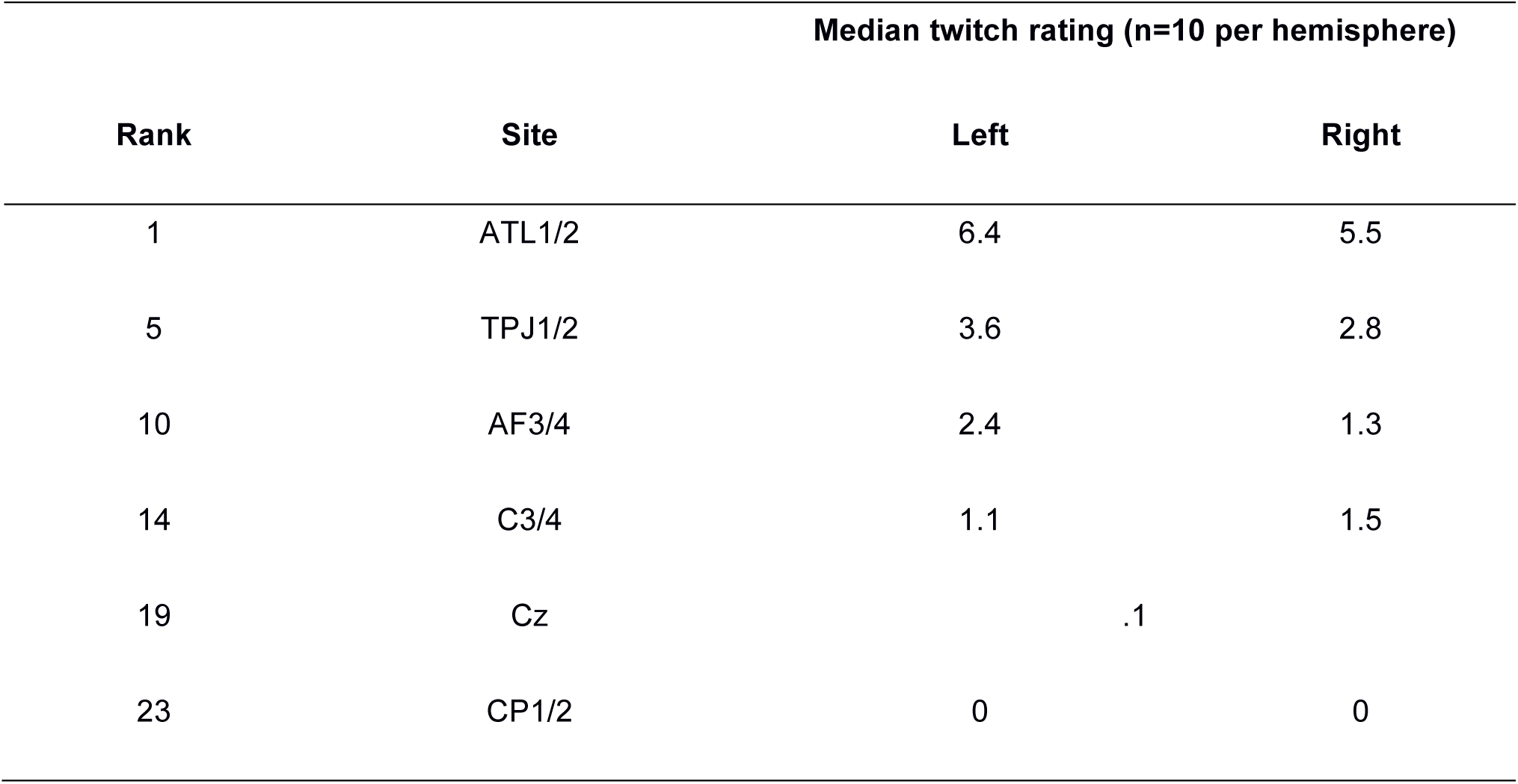
Sites chosen for control study of TMS intensity

| Rank | Site | Median twitch rating (n=10 per hemisphere) |  |
| --- | --- | --- | --- |
|  |  | Left | Right |
| 1 | ATL1/2 | 6.4 | 5.5 |
| 5 | TPJ1/2 | 3.6 | 2.8 |
| 10 | AF3/4 | 2.4 | 1.3 |
| 14 | C3/4 | 1.1 | 1.5 |
| 19 | Cz | .1 |  |
| 23 | CP1/2 | 0 | 0 |

One participant who had taken part in the main experiment was recruited. The participant received 5 single pulses of biphasic TMS at ∼0.2Hz at each location (in both left and right hemispheres for all sites except Cz), with the handle of the coil pointing South. After each set of 5 TMS pulses, the participant rated the subjective annoyance, pain, and twitches just as in the main experiment. An experimenter observed the participant, and noted down how many of the trials were accompanied by observable muscle twitches in the face, scalp, neck, or body of the participant.

After each set of 5 trials, the TMS coil was repositioned and the intensity of the TMS was changed according to a fully-randomised sequence. Five TMS intensities were used: 30, 40, 50, 60, and 70% of the maximum stimulator output. The mean ratings as a function of TMS intensity are shown in Figures C1 (mean across all 11 sites for 4 different ratings) and C2 (mean across left and right hemispheres for each site, excluding Cz).

### Results & Discussion

For each variable averaged across TMS sites, mean ratings increased linearly as a function of TMS intensity – r^2^s ranged between 0.90 (twitches) to 0.99 (visible twitches), all two-tailed ps<.05. The effect of TMS intensity was also approximately linear for ATL1/2 (r(3)=.912, p=.031), AF1/2 (r(3)=.982, p=.0029), TPJ1/2 (r(3)=.979, p=.0036, CP1/2 and Cz (both r(3)=.884, p=.047), but not for C3/4 (r(3)=.707, p=.18.

Data like this could be used, for example, to set TMS intensity for the control site to produce a comparable level of annoyance or muscle twitches to that of the target site (e.g., if control site cannot be varied freely). Further, these data support the assumption of a linear effect of TMS intensity on subjective annoyance, pain, and twitches, although for sites associated with low annoyance (midline and/or superior scalp locations), there may be an additional threshold of TMS intensity under which TMS is not at all annoying.

### Figures

**Figure C1:**
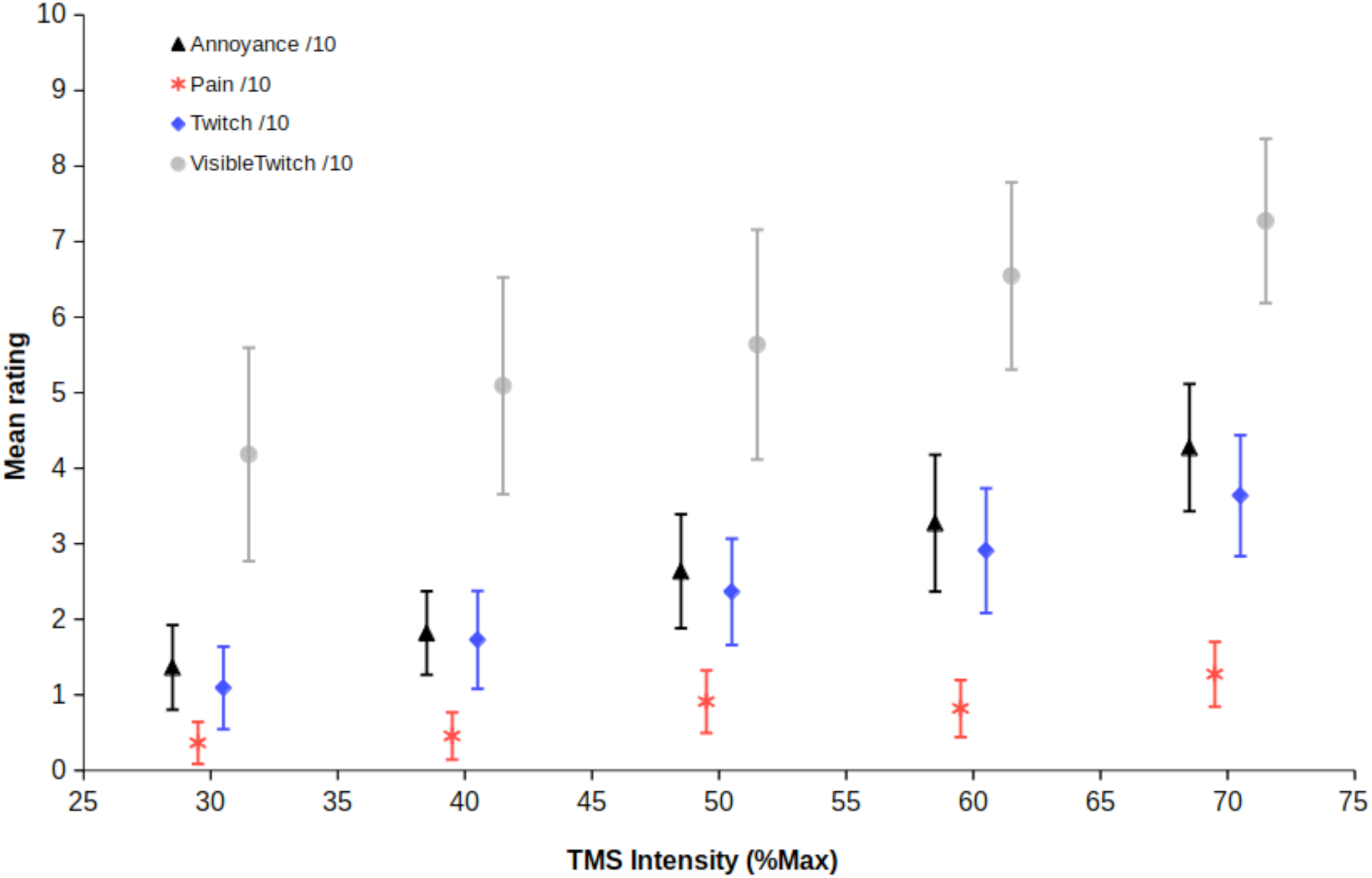
Mean across TMS sites as a function of TMS intensity (%Max) for subjective ratings of annoyance (black triangles), pain (red asterisks), and muscle twitches (blue diamonds), as well as observed twitches (grey circles). Visible twitches have been multiplied by two to show on the same scale.

**Figure C2:**
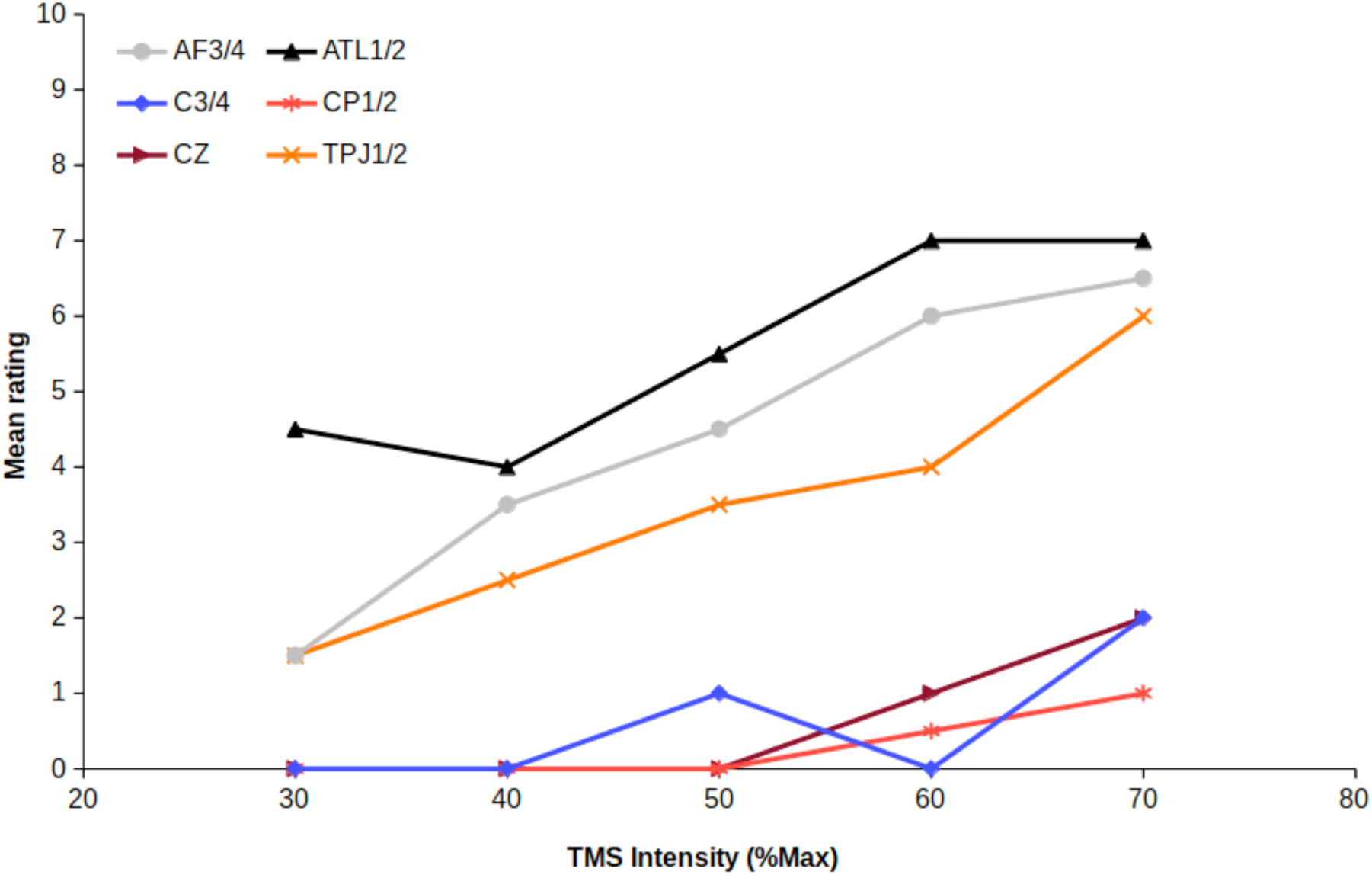
Mean across left and right hemispheres for each TMS site as a function of TMS intensity (%Max) for subjective ratings of annoyance. Black triangles: ATL1/2; Grey circles: AF3/4; Orange crosses: TPJ1/2; Blue diamonds: C3/4; Red asterisks: CP1/2; Maroon trianges: Cz.

## Appendix D / Supplementary materials 4: Scalp-to-brain distance for the 43 sites stimulated

### Methods

Twenty participants from an MRI dataset including the original TMS-SMART participant (NPH) were selected (10 females with a mean±SE age of 25.6±2.2 years; 10 males with a mean±SE age of 25.6±2.6 years).

Each participant’s anatomical MRI scan (MPRAGE, 1 × 1 × 1mm) was registered to NPH’s brain using 12 degree of freedom affine transformation in FLIRT (66). The resulting transform matrix was inverted using InvertXFM, then this inverted transform was applied to the 43 original locations used for TMS-SMART using ApplyXFM. This resulted, for each participant, in the approximate locations of scalp and brain sites targeted in the original experiment.

For each of 20 participants, the closest voxel on the scalp of the participant’s MRI to the transformed target location was estimated (x, y, and z coordinates in scanner anatomical space). From this scalp voxel, the closest voxel of grey matter (cortical or cerebellar) was estimated. The distance between scalp and grey matter was calculated, and plot in the images below.

### Results 1: Scalp to brain distances and correlations with Twitches and RTs

The mean±SE distance from scalp to brain across all 43 locations was 14.76±0.39 cm. The means and bootstrapped 95% confidence intervals are given in Table 1, main text.

**Figure D1.**
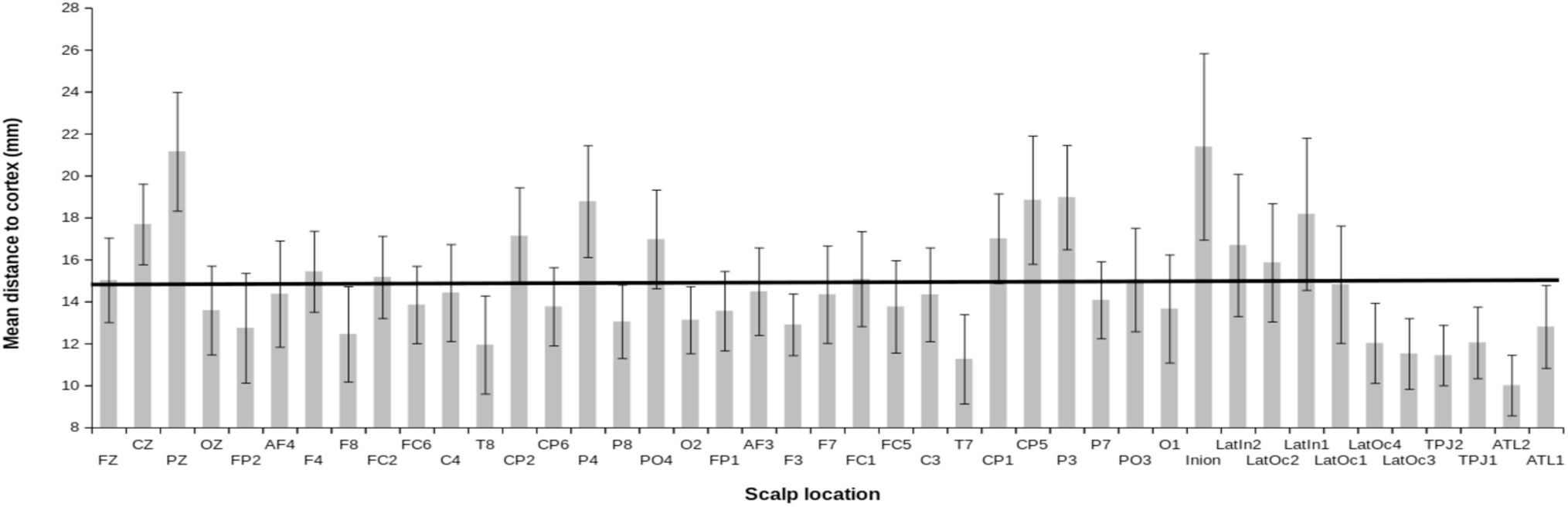
Mean scalp-to-brain distance for all 43 target locations. N=20. Solid horizontal line shows group mean. Error bars show 99.9% confidence intervals (i.e., p<.05, bonferroni corrected for 43 locations).

The mean across coil orientations of the median ratings of twitches (rated on a scale of 0- 10) from the main experiment, and the mean effect of TMS on reaction times (in seconds) were correlated across the locations targeted. These data were taken from a different groups of participants (i.e. those who participated in the original study).

These two variables were significantly correlated with scalp-to-brain distance. For twitches, there was a significant negative correlation (r(41)=-.525, p=.0003); as mean distance to cortex increases, ratings for twitches decrease (Figure D2). For the RT difference caused by TMS, there was a significant negative correlation (r(41)=-.400, p=.008); as mean distance to cortex increases, the RT difference under TMS decreases.

**Figure D2.**
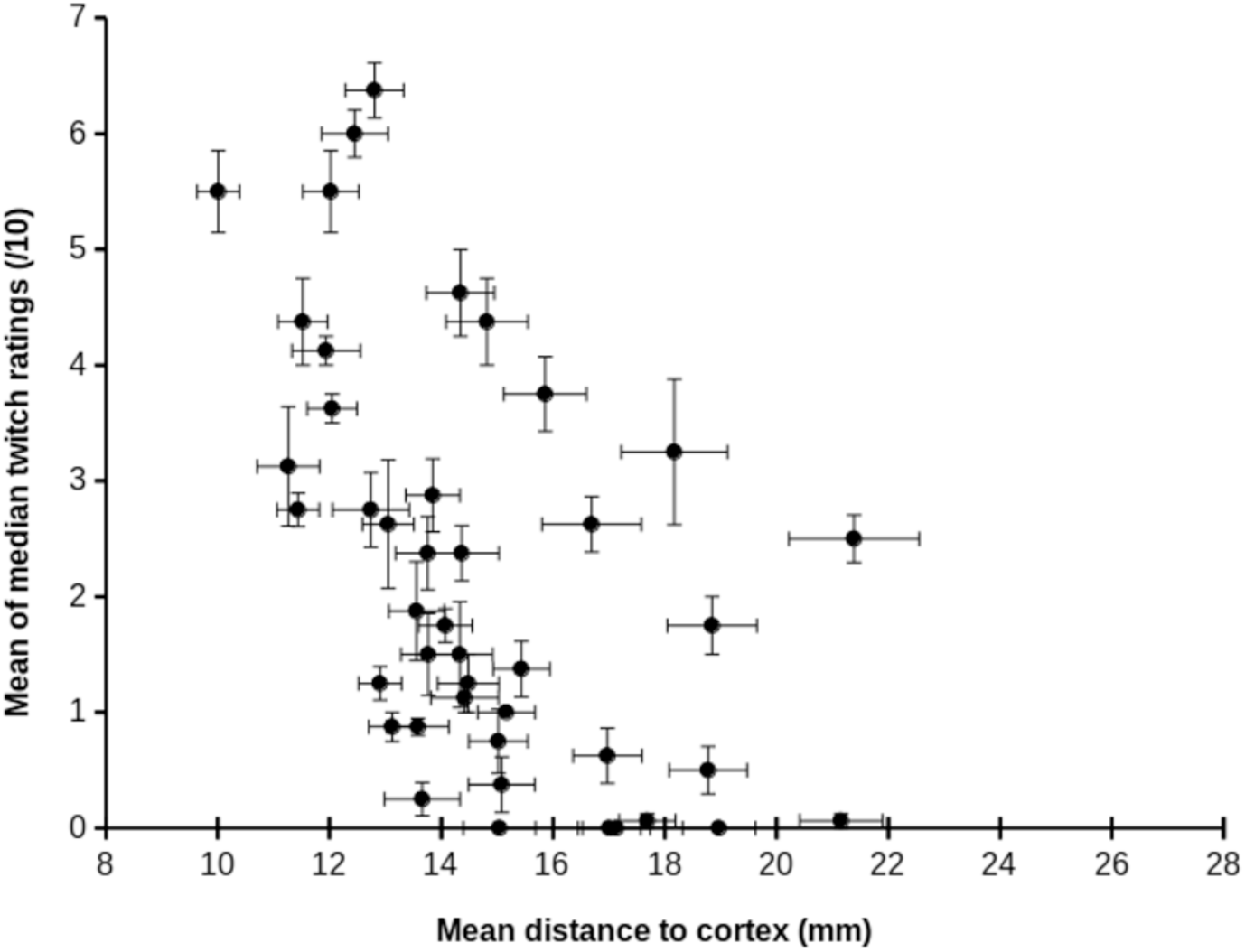
Mean±SE scalp-to-brain distances (mm) for 43 target locations (N=20) and mean±SE of median twitch ratings across the four coil orientations in the main experiment. Error bars show standard error. The relationship between the two variables was significant, r(41)=-.525, p=.0003.

**Figure D3.**
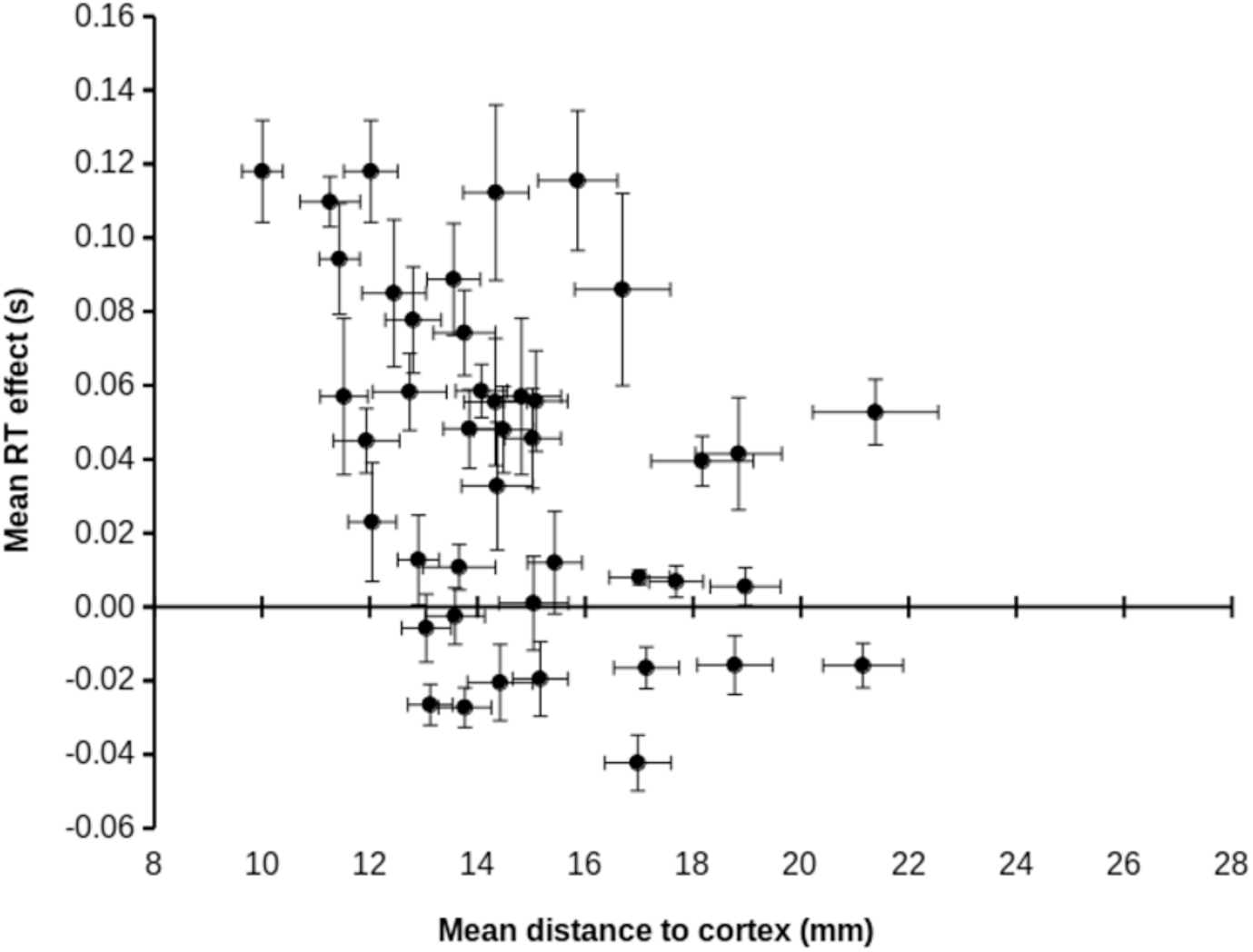
Mean±SE scalp-to-brain distances (mm) for 43 target locations (N=20) and mean±SE RT effects across the four coil orientations in the main experiment. Error bars show standard error. The relationship between the two variables was significant, r(41)=-.400, p=.008.

### Results 2: Scalp to brain distance as a model covariate

To check whether scalp to brain distance provides a better explanation for RT differences under TMS than subjective ratings of Twitches, we repeated our analyses (detailed in 3.3 and 3.4 main text). We replaced Scalp Location with Scalp to brain distance. It was not possible to enter Scalp Location and Scalp to brain distance together, as the two variables are perfectly collinear (i.e. there is one scalp to brain distance measure per location, so the two variables are perfectly predictable from each-other). For these models, Scalp to brain distance and Twitches were scaled to reduce collinearity.

The models replicated our previous finding. When Scalp to brain distance was entered alone, it was a significant predictor of RT differences under TMS (see Table D1; estimate = − 10.70, 95% CI = −18.52 – −2.88). As Scalp to brain distance decreases, the RT effects of TMS increase. Just as in our original models, Scalp Location accounted for significant variation in RT differences under TMS. The effect of Task was marginal, with the Flanker task having a trend for longer RTs (estimate = 34.87, 95% CI = −10.54 – 80.27).

**Table D1:**
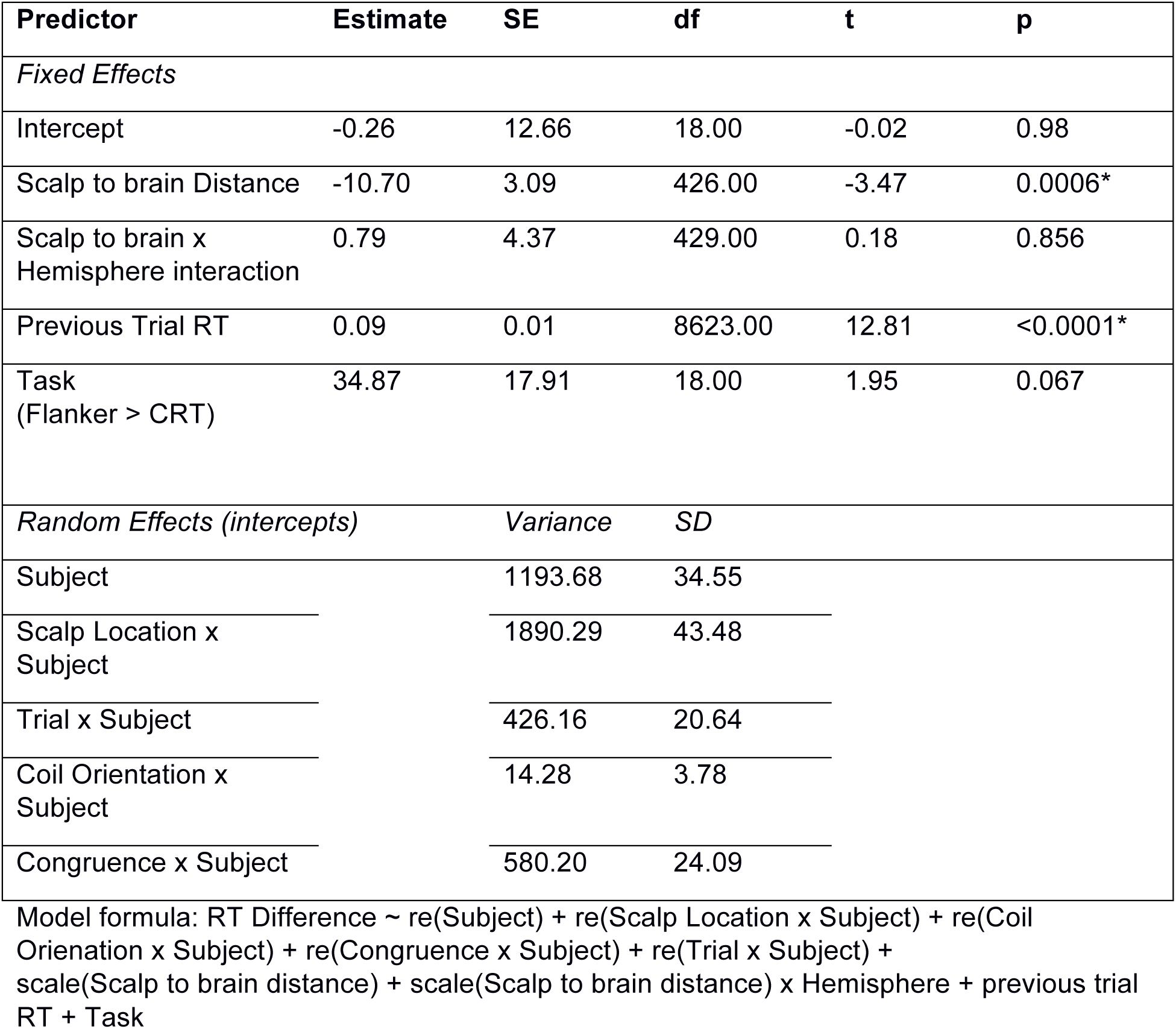
Model including Scalp-to-brain distance

We then added Twitch ratings to this model as a main effect and its interaction with Task (Table D2). In this model, Scalp to brain distance was no longer a significant predictor (estimate = 1.33, 95% CI = −6.10 – 8.76). Twitch ratings significantly predicted the RT difference under TMS (estimate = 20.10, 95% CI = 13.14 – 26.95), for every unit increase in Twitch ratings the RT cost of TMS increased by ∼20ms. The interaction of Twitch ratings and Task was also significant (estimate = 12.25, 95% CI = 2.87 – 21.63). A unit increase in Twitch ratings had a greater cost on the Flanker task, increasing RTs by ∼13ms more than for the CRT task (estimate = 12.25, 95% CI = 2.86 – 21.63).

**Table D2:** Model including Scalp-to-brain distance and Twitch ratings

| Predictor | Estimate | SE | df | t | p |
| --- | --- | --- | --- | --- | --- |
| *Fixed Effects* |  |  |  |  |  |
| Intercept | -0.35 | 13.10 | 18.00 | -0.03 | 0.98 |
| Scalp to brain Distance | 1.33 | 2.80 | 436.00 | 0.48 | 0.64 |
| Scalp to brain x Hemisphere interaction | -0.35 | 3.77 | 422.00 | -0.09 | 0.93 |
| Previous Trial RT | 0.09 | 0.01 | 8643 | 12.52 | <0.0001* |
| Twitch Ratings | 20.10 | 2.60 | 632.00 | 7.71 | <0.0001* |
| Twitch Ratings x Task interaction (Flanker > CRT) | 12.25 | 3.54 | 605.00 | 3.46 | 0.0006* |
| *Random Effects (intercepts)* |  | *Variance* | *SD* |  |  |
| Subject |  | 1327.26 | 36.43 |  |  |
| Scalp Location x Subject |  | 1309.03 | 36.18 |  |  |
| Trial x Subject |  | 434.05 | 20.83 |  |  |
| Coil Orientation x Subject |  | 15.71 | 3.96 |  |  |
| Congruence x Subject |  | 581.26 | 24.11 |  |  |

Model formula: RT Difference ~ re(Subject) + re(Scalp Location x Subject) + re(Coil Orientation x Subject) + re(Congruence x Subject) + re(Trial x Subject) + scale(Scalp to brain distance) + scale(Scalp to brain distance) x Hemisphere + previous trial RT + scale(Twitch ratings) + Task x scale(Twitch ratings)

### Conclusion

We found that Scalp to brain distance was significantly correlated with both subjective ratings of twitches and RT differences under TMS. This is likely a product of physiology – areas of the brain closest to the scalp (frontal and inferior sites) also have a greater density of muscles and nerve fibres; this leads to greater discomfort. However, in models where both are taken into account, subjective discomfort is the stronger predictor of RT costs under TMS. This supports our original analysis (see 3.3 and 3.4 in the main text).

